# Interleukin-1β Drives Disease Progression in Arrhythmogenic Cardiomyopathy

**DOI:** 10.1101/2024.12.11.628020

**Authors:** Vinay R. Penna, Junedh M. Amrute, Morgan Engel, Emily A. Shiel, Waleed Farra, Elisa N. Cannon, Colleen Leu-Turner, Pan Ma, Ana Villanueva, Haewon Shin, Alekhya Parvathaneni, Joanna Jager, Carlos Bueno-Beti, Angeliki Asimaki, Kory J. Lavine, Jeffrey E. Saffitz, Stephen P. Chelko

## Abstract

Arrhythmogenic cardiomyopathy (ACM) is a genetic form of heart failure that affects 1 in 5000 people globally and is caused by mutations in cardiac desmosomal proteins including *PKP2, DSP*, and *DSG2.* Individuals with ACM suffer from ventricular arrhythmias, sudden cardiac death, and heart failure. There are few effective treatments and heart transplantation remains the best option for many affected individuals. Here we performed single nucleus RNA sequencing (snRNAseq) and spatial transcriptomics on myocardial samples from patients with ACM and control donors. We identified disease-associated spatial niches characterized by co-existence of fibrotic and inflammatory cell types and failing cardiac myocytes. The inflammatory-fibrotic niche co-localized to areas of cardiac myocyte loss and was comprised of *FAP* (fibroblast activation protein) and *POSTN* (periostin) expressing fibroblasts and macrophages expressing *NLRP3* (NLR family pyrin domain containing 3) and NFĸB activated genes. Using homozygous Desmoglein-2 mutant (*Dsg2^mut/mut^*) mice, we identified analogous populations of *Postn* expressing fibroblasts and inflammatory macrophage populations that co-localized within diseased areas. Detailed single cell RNA sequencing analysis of inflammatory macrophage subsets that were increased in ACM samples revealed high levels of interleukin-1β (*Il1b*) expression. To delineate the possible benefit of targeting IL-1β in ACM, we treated *Dsg2^mut/mut^* mice with an anti-IL-1β neutralizing antibody and observed attenuated fibrosis, reduced levels of inflammatory cytokines and chemokines, preserved cardiac function, and diminished conduction slowing and automaticity, key mechanisms of arrhythmogenesis. These results suggest that currently approved therapeutics that target IL-1β or IL-1 signaling may improve outcomes for patients with ACM.

## Introduction

Arrhythmogenic cardiomyopathy (ACM) is a familial non-ischemic heart disease, affecting 1:2,000 to 1:5,000 people, globally *(1)*. It is one of the leading causes of sudden cardiac death (SCD) in young individuals due to malignant ventricular arrhythmias *(1, 2)*. Individuals with ACM may also progress to end-stage heart failure. Currently, treatments such as antiarrhythmics, standard medical therapies for heart failure, and implantable cardioverter defibrillators (ICDs) only temporize symptoms, and heart transplantation represents the only curative therapy *(3, 4)*.

ACM is primarily caused by mutations in cardiac desmosomal genes *(5).* While other cardiac cytoskeletal and ion transport genes may present with an ACM-like phenotype, most ACM cases stem from mutations in the desmosomal genes plakophilin-2 (*PKP2*), desmoplakin (*DSP*), and desmoglein-2 (*DSG2*) *(5).* How these mutations result in ACM is not clearly understood, but it is thought to involve abnormal NFκB, Wnt/β-catenin, and Hippo signaling pathways *(6–8)*. Recent studies have recognized that ACM is associated with a striking cardiac and systemic inflammatory response *(9)*.

Prominent myocardial fibrosis and inflammation have been observed in over 70 percent of autopsy samples from patients with ACM. Furthermore, patients with ACM display elevated serum levels of inflammatory cytokines *(1, 2, 9, 10)*. We have previously reported that signaling mediated via NFκB, a master regulator of the innate immune response, is activated in a mouse model of ACM harboring homozygous knock-in of a variant in the gene encoding the desmosomal protein, desmoglein-2 (*Dsg2^mut/mut^* mice) *(11)*. We observed that using a genetic approach to block NFκB signaling in cardiac myocytes alone is sufficient to prevent myocardial loss and fibrosis, preserve contractile function, and suppress arrhythmias in *Dsg2^mut/mut^* mice *(12)*. We also found that NFκB signaling in cardiac myocytes leads to a 5-fold increase in monocytes and macrophages expressing C-C motif chemokine receptor-2 (CCR2), a potent chemotactic molecule that has been implicated in adverse cardiac remodeling and fibrosis *(13–15)*. Suppression of CCR2+ monocyte and macrophage recruitment to the heart was sufficient to halt progression of ACM pathology in *Dsg2^mut/mut^*mice *(12)*.

Despite these recent insights, significant gaps remain in our understanding of how inflammatory monocyte and macrophage populations contribute to heart failure, myocardial inflammation and fibrosis, and arrhythmogenesis. Little is known regarding the cellular composition of ACM lesions or the key mediators of cardiac inflammation. Improved understanding of the cellular and transcriptomic landscape of ACM lesions and the aberrant cell signaling pathways utilized to drive tissue pathology will be critical to identify new therapeutic targets for this devastating disease.

To address these gaps in knowledge, we performed single nucleus RNA sequencing (snRNAseq) on myocardial samples from patients with clinically active ACM (n=6, 3 patients with *DSP* variants and 3 patients with *PKP2* variants) and donor controls (n=12, no history of heart disease). In addition, we performed spatial transcriptomic sequencing on ACM patient samples (n=3, 2 patients with *PKP2* variants and 1 patient with a *DSP* variant) and donor controls (n=2). Using these data, we deconvoluted the cellular landscape of ACM, identified ACM-associated disease signatures, and uncovered spatially restricted niches containing pro-fibrotic fibroblasts and inflammatory macrophages that localized to areas of myocardial disease. Using an established mouse model of ACM (i.e., *Dsg2^mut/mut^* mice), we observed analogous cell populations and niches with enriched expression of inflammatory mediators including interleukin-1β (IL-1β). To establish a causative relationship between IL-1 signaling and cardiac pathology, we treated *Dsg2^mut/mut^*mice with a neutralizing antibody against IL-1β and observed significant improvements in myocardial pathology and function. These findings highlight a role for targeting IL-1 signaling in ACM.

## Results

### Single nucleus RNA (snRNAseq) sequencing reveals the cellular landscape of ACM

We performed snRNAseq on transmural left ventricular specimens obtained from the apical anterior left ventricular wall of donor controls (n=12) and patients with ACM (n=6, including 3 patients with a *DSP* variant and 3 with a *PKP2* variant) (table S1) at the time of heart transplantation (**Fig. 1A**). Following doublet removal and quality control (fig. S1A), we performed dimensional reduction, UMAP construction, and differential gene expression to annotate cell types (**Fig. 1B**). We identified 14 transcriptionally distinct cell types expressing canonical marker genes (**Fig. 1C**). In addition, we constructed cell type specific gene set scores and detected strong separation across clusters (fig. S1B). Analysis of cell type composition demonstrated a robust expansion of fibroblast, myeloid, and T-cell populations in ACM myocardium compared to donor controls (**Fig. 1D**). To identify how ACM pathogenic variants perturb the transcriptional profile in a cell type specific manner, we performed pseudobulk differential expression analysis at the patient level and tabulated up- and down-regulated genes in ACM samples compared to donor controls. We found fibroblasts, endothelial cells, cardiac myocytes, pericytes, endocardial cells, and myeloid cells harbored the most prominent transcriptional changes (**Fig. 1, E and F**). These data suggest that ACM is associated with remodeling of major cell types including immune and stromal components of the myocardium.

**Fig 1.**
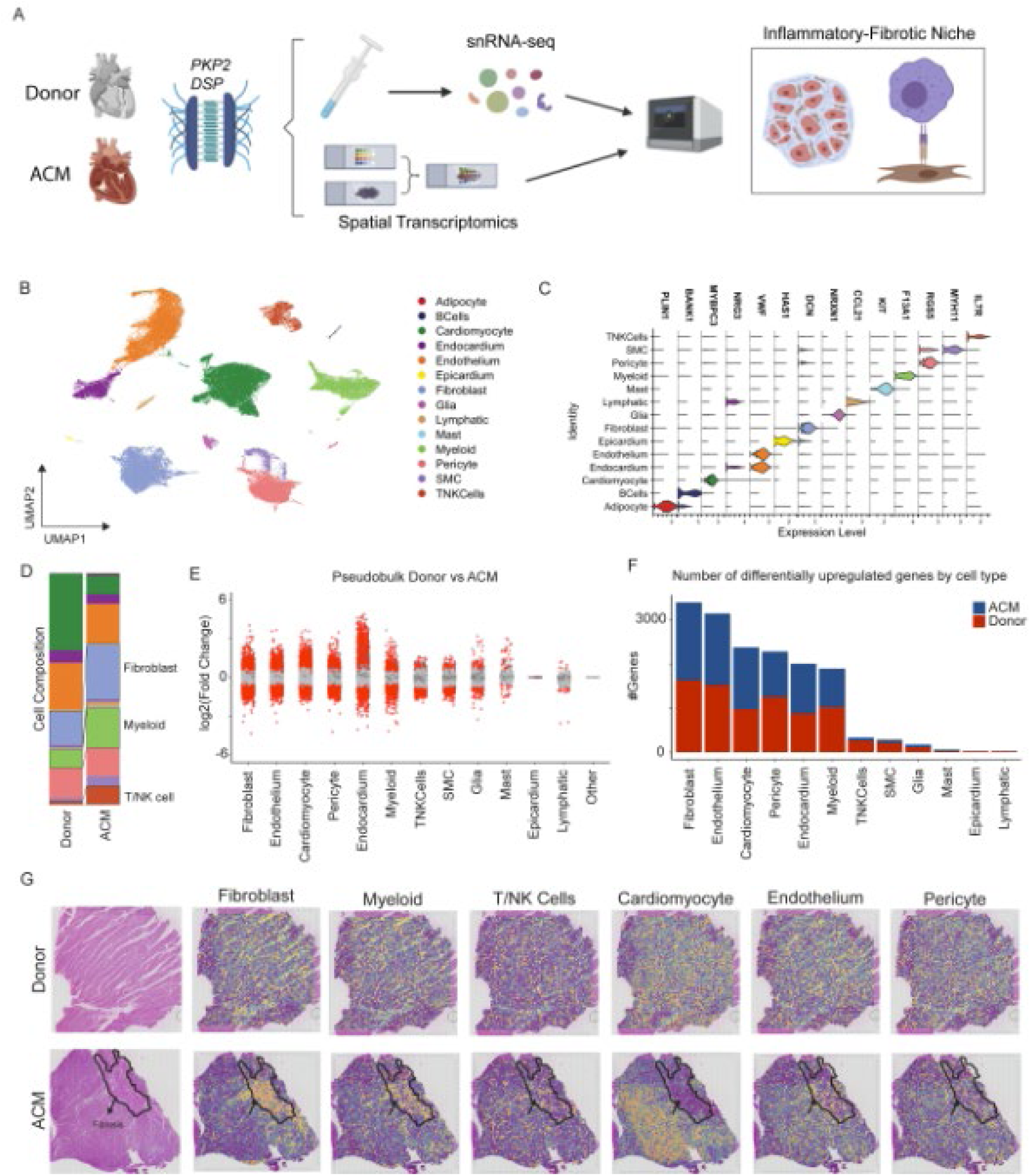
ACM alters the cardiac cellular and transcriptomic environment. (**A**) Study design schematic outlining human tissue sequencing methods. (**B**) Global UMAP with annotations of major cell populations. (**C**) Violin plot outlining major canonical markers identifying cell populations. (**D**) Composition plot displaying relative proportions of major cell types between donor control group and ACM group. (**E**) Pseudobulk analysis displaying degree of gene expression changes across major cell types in Donor vs ACM. (**F**) Number of total differentially upregulated genes for each cell type. (**G)** Spatial transcriptomic plots displaying major cell populations overlaid over H&E tissue images.

### Spatial transcriptomics reveals formation of an inflammatory-fibrotic niche in ACM

To define the spatial organization of cell types enriched in ACM, we performed spatial transcriptomics sequencing of donor control (n=2) and ACM (n=3) left ventricular tissues. H&E staining showed areas of disrupted myocardial architecture and fibrosis, which we refer to as ACM lesions (fig. S2). ACM specimens including those with pathogenic *PKP2* (n=2) and *DSP* (n=1) variant (table S1, **Fig. 1A**). Each processed sample displayed high quality UMI counts *(16, 17)* (fig S3). As the spatial resolution of formalin-fixed, paraffin-embedded (FFPE) Visium technology is ∼40 μm^2^, each spot contains numerous cell types. To decipher the proportion of different cell types within each spot, we applied Tangram, which leverages our paired spatial transcriptomic and snRNAseq data sets to map cells into space *(18)*. To visualize localization of major cell types, we plotted deconvolution scores for the major cell types overlaid on the H&E-stained image (**Fig. 1G**). In donor hearts, cardiac myocytes represented the dominant cell population with homogenous organization of macrophages and fibroblasts. In ACM samples, we observed areas depleted of cardiac myocytes that were enriched with macrophages and fibroblasts, which corresponded with ACM lesions. Endothelial cells, pericytes, and T-cells were homogenously distributed across the myocardium (**Fig. 1G**).

To further characterize the spatial architecture of ACM, we independently clustered the aggregated donor and ACM spatial transcriptomic datasets to identify unique spatial niches. Unbiased clustering identified 7 transcriptionally distinct spatial niches (**Fig. 2A**). Niches 0 and 1 were enriched in donor compared to ACM samples and contained cardiac myocytes, macrophages expressing tissue resident markers, and fibroblasts. Niches 2 and 3 were enriched in ACM relative to donor controls and contained cardiac myocytes expressing heart failure markers (*NPPA*, *NPPB*, and *ANKRD1*) *(19–22)*. Niches 4, 5, and 6 were also increased in ACM relative to donor controls and were composed of inflammatory macrophages and pro-fibrotic fibroblasts (**Fig. 2, A and B**). We then co-registered the niche assignments onto the H&E-stained images and found that Niche 4 was highly enriched in ACM lesions (**Fig. 2C**).

**Fig 2.**
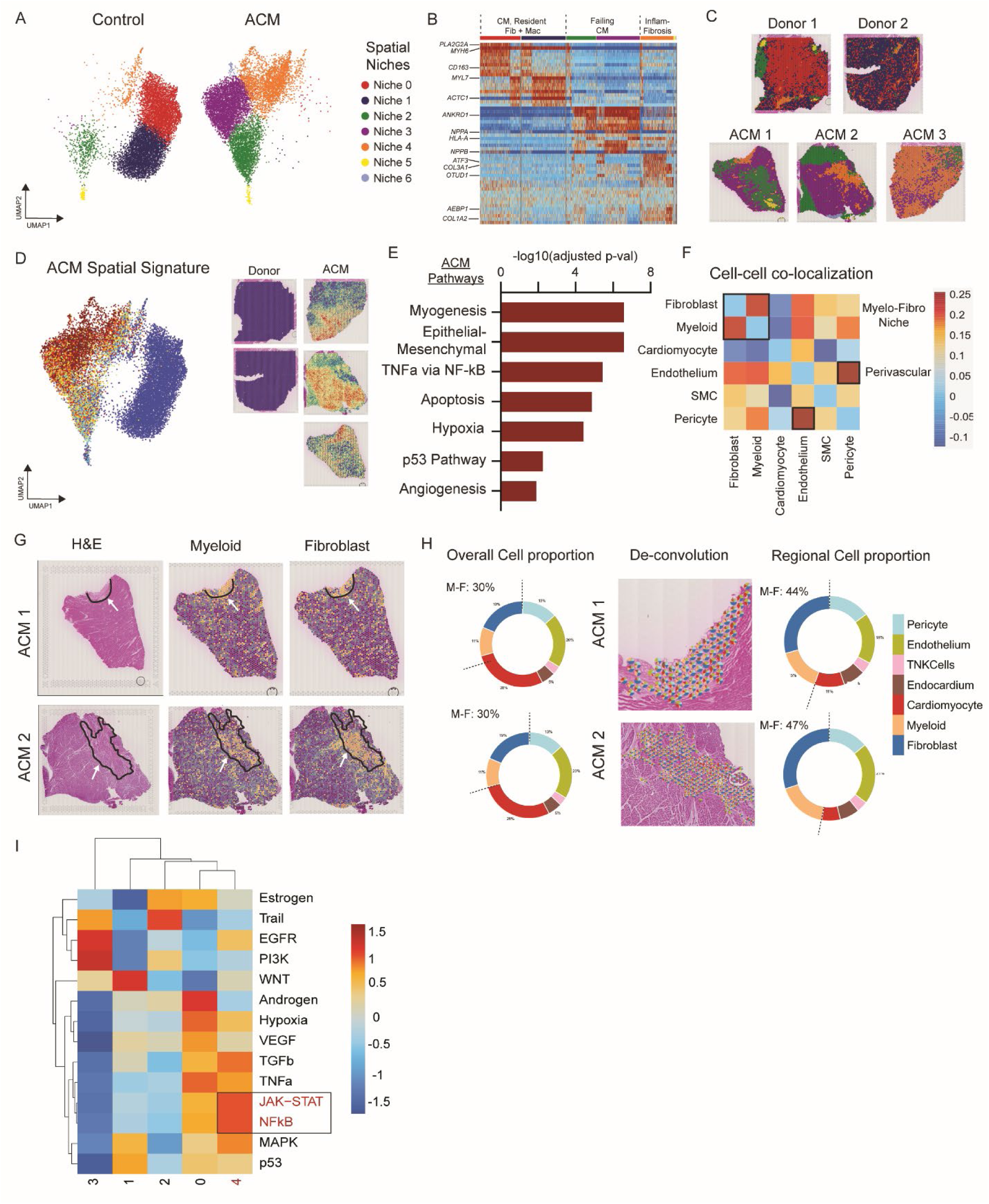
ACM has a unique spatial niche, which includes strong overlap between myeloid cells and fibroblast. (**A**) Global UMAPs derived from spatial gene expression data outlining major niches. (**B)** Heatmap displaying genes expressed by spatial niches. (**C**) Spatial niches overlaid onto tissue architecture to determine where niches are positioned in tissue space. (**D**) Top differentially expressed genes between ACM and Donor controls overlaid onto the spatial UMAP. (**E**) Pathway analysis displaying major upregulated pathways in ACM relative to Donor controls. (**F**) Pearson correlation plot displaying likelihood that two cell types will be found in the same spatial location in tissue. Darker blue indicates higher probability. (**G**) H&E images from two ACM samples highlighting areas of tissue damage, myeloid, and fibroblast concentration. (**H**) Circle graph displaying overall cell type proportions in tissue along with a region-specific cell type proportion graph in areas of tissue damage and fibrosis. (**I**) Heatmap displaying expression of major signaling pathways across spatial niches.

To identify genes enriched in ACM associated spatial niches, we performed a differential expression analysis between ACM and donor control samples and overlaid the signature onto the UMAP embedding (**Fig. 2D**). This analysis revealed enrichment of the ACM signature in Niche 3, 4, and 6 with strong co-localization to areas within and surrounding ACM lesions. Pathway enrichment analysis showed that the ACM transcriptional signature was enriched for myogenesis, epithelial-mesenchymal transition, TNFα via NFĸB, apoptosis, hypoxia, p53, and angiogenesis terms (**Fig. 2E**).

To further characterize the ACM niches and identify which cell types co-localize in these spaces, we built a Pearson correlation coefficient from tangram deconvolution scores and found that macrophages and fibroblasts form a myelo-fibro niche while endothelial cells and pericytes form a perivascular niche (**Fig. 2F**). We then overlaid tangram deconvolution scores for macrophages and fibroblasts in multiple ACM samples and saw strong co-localization of macrophage/fibroblast scores with areas of fibrosis (**Fig. 2G**). To characterize relative cell abundance, we constructed aggregate pie charts across the entire tissue and found macrophages and fibroblasts were quantitatively expanded in ACM lesions (**Fig. 2, G and H**). To infer active signaling events in ACM lesions, we used PROGENy for pathway analysis and discovered marked enrichment for NFĸB and JAK-STAT signaling in Niche 4 (the inflammatory-fibrotic niche) (**Fig. 2I**).

### snRNAseq reveals expansion of POSTN expressing fibroblasts and inflammatory macrophages in ACM

Given the enrichment of macrophages and fibroblasts within ACM lesions, we sought to characterize their precise cell states. To dissect the heterogeneity of fibroblasts and macrophages, we performed unbiased clustering of these populations. We identified 7 transcriptionally distinct fibroblast cell states (Fib 1-7): Fib1 (*ACSM3, APOD*), Fib2 (*KAZN, LSAMP*), Fib3 (*POSTN, THBS4*), Fib4 (*PCOLCE2, PDZRN4*), Fib5 *(CCDC80, COL15A1*), Fib6 (*TNC, RUNX1*), and Fib7 (*FOSB, FOS*) (fig S4A, **Fig. 3A**). Fib1 and Fib2 were enriched in donor controls, while Fib3 and Fib7 were enriched in ACM samples (**Fig. 3B**). We calculated genes differentially expressed in donor control and ACM samples and visualized their expression using density plots. Genes with highest enrichment in donor control fibroblasts (*ACSM3)* were expressed in Fib 1, while genes enriched in ACM (*THBS4, RUNX1*, and *POSTN*) were predominately expressed in Fib 3 (**Fig. 3C**). To identify spatial niches enriched with pro-fibrotic cell states we plotted *ACTA2*, *THBS4*, *FAP*, *POSTN*, *COL1A1,* and *RUNX1* (genes that have been previously implicated in tissue fibrosis and pathological remodeling *(19, 23)* across the spatial niches and found maximal enrichment in Niche 4 (**Fig. 3D**). Consistent with the above analysis, we performed pseudobulk differentially gene expression in fibroblasts using our snRNAseq data and found that the donor control and ACM fibroblast signatures were enriched in Fib 1-2 and Fib 3, 6-7, respectively (**Fig. 3E**). Moreover, the ACM fibroblast signature co-localized with Niches 3-4, areas corresponding to and surrounding ACM lesions (**Fig. 3F**).

**Fig 3.**
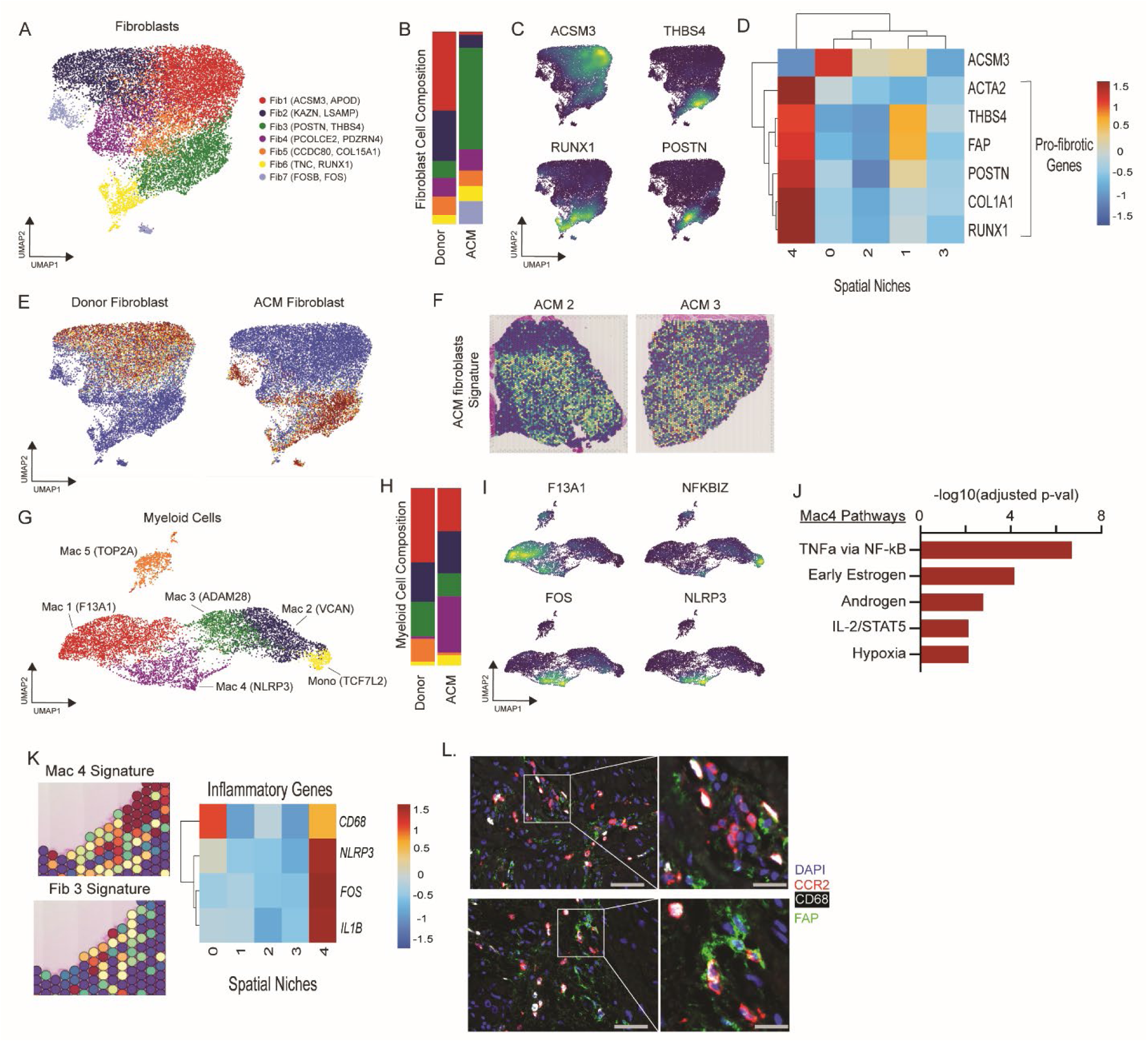
POSTN+ fibroblasts and inflammatory macrophages are increased in ACM and colocalize in areas of tissue damage and fibrosis. (**A**) UMAP of fibroblast populations. (**B**) Composition plots of fibroblast populations between Donor controls and ACM. (**C**) Major fibroblast gene markers overlaid on the fibroblast UMAP. (**D**) Heatmap displaying expression of fibroblast markers across spatial niches. (**E**) Fibroblast differential gene expression signature associated with either Donor controls or ACM overlaid onto the fibroblast UMAP space. (**F**) ACM fibroblast gene expression signature overlaid onto ACM tissue space. (**G**) UMAP of myeloid populations. (**H**) Composition plots of myeloid populations between Donor controls and ACM. (**I**) Major gene markers for myeloid populations overlaid on the myeloid UMAP. (**J**) Pathway analysis displaying pathways upregulated in ACM relative to Donor controls based on differentially expressed myeloid genes. (**K**) Colocalization of the inflammatory macrophage population (Mac4) and the POSTN+ fibroblast population (Fib3) in areas of tissue damage and fibrosis and heatmap displaying expression of major inflammatory genes across spatial niches. (**L**) Immunofluorescence staining displaying colocalization of CCR2+ CD68+ macrophages and FAP+ fibroblasts in ACM tissue samples. Two independent samples were used. Broader images are captured at 20x.

We next sought to delineate macrophage states in ACM lesions. Unbiased clustering of myeloid cells within our snRNAseq data revealed 6 distinct cell states: Monocytes (*TCF7L2*), Mac1 (*F13A1*), Mac2 (*VCAN*), Mac3 (*ADAM28*), Mac4 (*NLRP3*), and Mac5 (proliferating, *TOP2A*) (fig S4B, **Fig. 3G**). Cell state composition analysis showed expansion of proliferating cells in donor controls (consistent with prior studies showing decreased myeloid proliferation in genetic DCM) *(19, 24, 25)* and expansion of *NLRP3*+ pro-inflammatory macrophages (Mac4) and monocytes in ACM (**Fig. 3, G and H**), consistent with prior a report *(26)*. Expression of *F13A1* localizes to the resident macrophage population while *FOS* and *NLRP3* are enriched in Mac4, and *NFKBIZ* is enriched in monocytes (**Fig. 3I**). Pathway analysis of the marker genes for Mac4 show enrichment for TNFα via NFĸB, early estrogen, androgen, IL-2/STAT5, and hypoxia activation (**Fig. 3J**). These findings highlight that in ACM inflammatory monocytes and macrophages show enriched NFĸB-dependent signaling. We then plotted the gene signature of Mac4 and Fib3 in an ACM sample focusing on the regions of fibrosis and found a strong overlap between inflammatory macrophages and pro-fibrotic fibroblasts. Additionally, we plotted inflammatory genes such as *NLRP3, FOS, and IL1B,* which have previously shown to be enriched in inflammatory macrophages *(13–15, 19, 23, 27)* and found enrichment of these genes in spatial Niche 4. Collectively, these findings support the idea that inflammatory macrophages and pro-fibrotic fibroblast cell states are enriched in ACM lesions and may signal to one another, serving as a driving factor in the development of fibrosis in ACM, as demonstrated previously in myocardial infarction *(23, 28)* (**Fig. 3K**). To validate predictions of cell composition within ACM lesions, we performed immunofluorescence staining for inflammatory macrophages (CD68+ and CCR2+) and activated fibroblasts (FAP+) and observed co-localization of both populations in ACM lesions (**Fig. 3L**).

### snRNAseq of *Dsg2*^mut/mut^ mice reveals expansion of analogous pro-fibrotic fibroblasts and inflammatory macrophages

To explore the contribution of macrophages and fibroblast populations in ACM pathogenesis, we utilized the homozygous Desmoglein-2 mutant (*Dsg2^mut/mut^*) mouse model of ACM. This strain recapitulates major pathological and physiological characteristics of ACM, including inflammation, fibrosis, impaired cardiac function, and arrhythmias *(11, 12, 29)*. To determine if an analogous macrophage-fibroblast axis contributed to disease in *Dsg2^mut/mut^*mice, we performed targeted single cell RNA sequencing of the fibroblast and myeloid populations in 6-week-old *Dsg2^mut/mut^*mice and age-matched wild-type (WT) hearts (**Fig. 4A**). After performing quality control (fig S5A), murine fibroblast (mFib1-7) clustering identified 7 transcriptionally different clusters: mFib1 (*Morrbid, Pla1a)*, mFib2 (*Ccl19, L3mbtl4*), mFib3 (*Postn, Comp*), mFib4 (*Igfbp3, Cytl1*), mFib5 (*Opcml, Igfbp5*), mFib6 (*Cxcl14, Penk)*, and mFib7 (*Ptx3, Ccl2*) (fig S5B, **Fig. 4B**). mFib3 and mFib7 were enriched in *Dsg2^mut/mut^* mice, while mFib1, mFib5, and mFib6 were enriched in WT mice (**Fig. 4C**). To determine if the *Postn* enriched cluster in mice (mFib2) was similar to the *POSTN* enriched cluster in humans (Fib3), we generated a gene set signature score using the human genetic expression data (converted into mouse orthologs) (table S2) and overlaid that score onto the mouse UMAP (**Fig. 4D**). The ACM human fibroblast gene signature was robustly expressed by the mFib2 cluster, suggesting the existence of a transcriptionally analogous population of fibroblasts present in *Dsg2^mut/mut^*mice (**Fig. 4E**).

**Fig 4.**
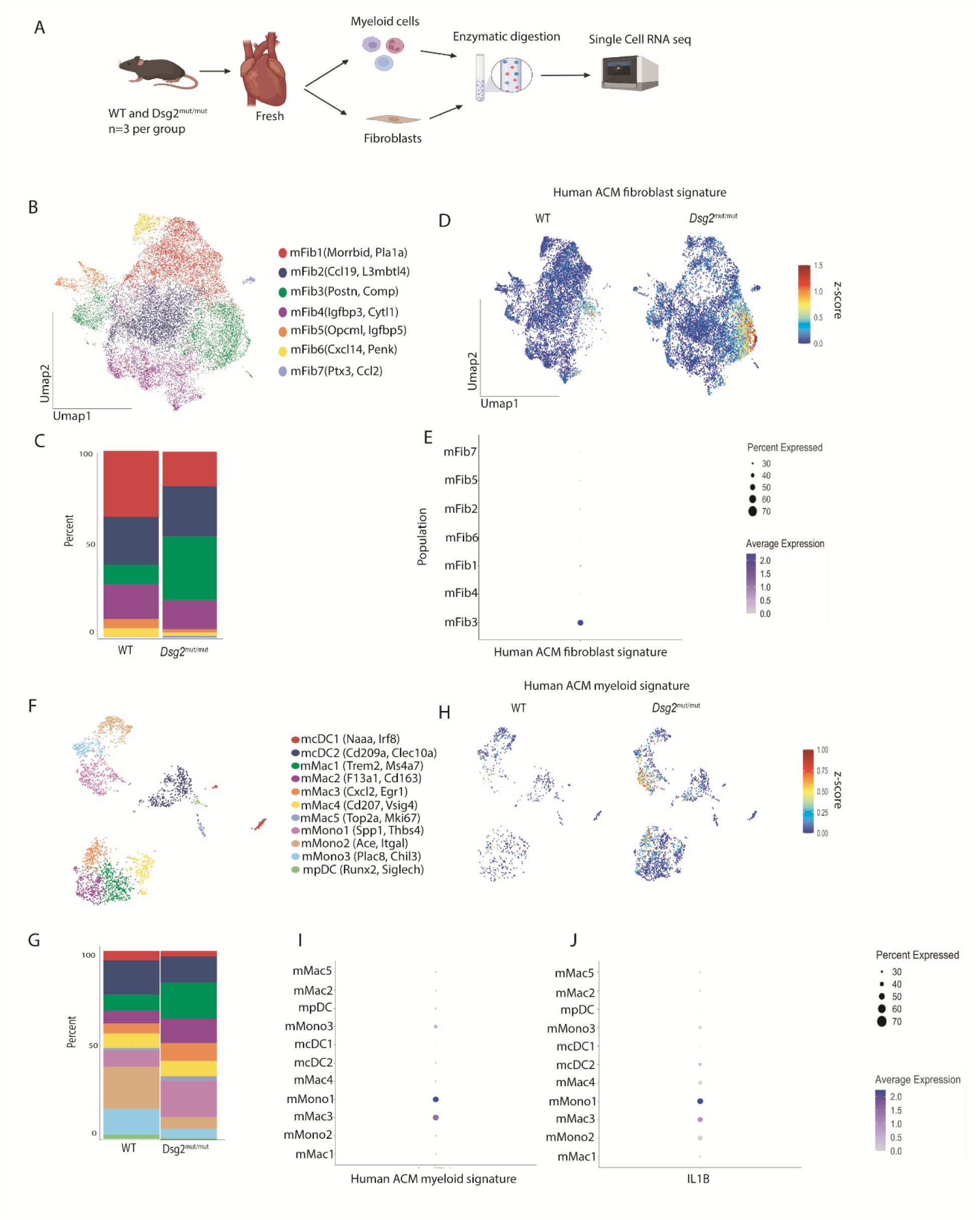
*Dsg2^mut/mut^* mice have analogous POSTN+ fibroblast and inflammatory macrophage populations as found in human ACM. (**A**) Study design outlining how myeloid and fibroblast libraries were sequenced. (**B**) Global UMAP of mouse fibroblast populations. (**C**) Composition plots comparing fibroblast populations between WT mice and *Dsg2^mut/mut^* mice. (**D**) Gene expression score generated from mouse orthologs of differentially upregulated fibroblast genes from human ACM sequencing data overlaid onto the mouse fibroblast UMAP. (**E**) The previously generated human gene expression score represented on a dot plot across mouse fibroblast populations. (**F**) Global UMAP of mouse myeloid populations. (**G**) Composition plots comparing myeloid populations between WT mice and *Dsg2^mut/mut^* mice. (**H**) Gene expression score generated from mouse orthologs of differentially upregulated myeloid genes from human ACM sequencing data overlaid onto the mouse myeloid UMAP. (**I**) The previously generated human gene expression score represented on a dot plot across mouse myeloid populations. (**J**) Expression score for *Il1b* measured across mouse myeloid populations.

Further analysis of the myeloid populations yielded 12 transcriptionally unique states: mcDC1 (*Irf8, Naaa*), mcDC2 (*Cd209a, Clec10a*), mMac1 (*Trem2, Ms4a7*), mMac2 (*F13a1, Cd163),* mMac3 (*Cxcl2, Egr1),* mMac4 (*Cd207, Vsig4*), mMac5 (*Top2a, Mki67),* mMono1 (*Spp1, Thbs4),* mMono2 (*Ace, Itgal)*, mMono3 (*Plac8, Chil3)*, and mPDC (*Runx2, Siglech*) (fig S5C, **Fig. 4F**). mMono1, mMac1, and mMac3 were significantly overrepresented in *Dsg2^mut/^*^mut^ mice compared to WT mice, while mMono2 and mMono3 were enriched in WT (**Fig. 4G**). Similar to the fibroblasts, we generated a gene set signature score using the human expression data (table S3) and overlaid it onto the mouse UMAP to determine if there was an analogous population of inflammatory macrophages in *Dsg2^mut/^*^mut^ mouse hearts (**Fig. 4H**). The ACM human myeloid signature was robustly expressed by mMono1 and mMac3 (**Fig. 4I**). Subsequent analysis revealed that these populations also robustly express *Il1b* (**Fig. 4J**), suggesting that these are an analogous population of inflammatory macrophages similar to those found in hearts from patients with ACM. Overall, these findings point to the presence of a similar inflammatory-fibroblast axis in *Dsg2^mut/mut^* mouse hearts, suggesting that this mouse model mirrors human ACM disease at the cellular and transcriptional level.

### Targeting IL-1β attenuates disease characteristics in *Dsg2^mut/mut^* mice

IL-1β is a primordial inflammatory cytokine of the innate immune response, produced predominantly by macrophages *(23)*. Recent studies, including the CANTOS and VCU-ART trials, have explored the efficacy of IL-1β blockade in various forms of cardiovascular disease including myocardial infarction and atherosclerosis *(30–33)*. Yet, the role of targeting IL-1 signaling in ACM is poorly understood. Given our findings of increased *Il1b* expressing inflammatory macrophages in *Dsg2*^mut/mut^ mouse hearts and enrichment for *NLRP3* in corresponding human macrophages, we set out to determine if IL-1β blockade can mitigate ACM disease progression prior to overt cardiac remodeling and fibrosis. Therefore, 8-week-old WT and *Dsg2*^mut/mut^ mice were treated with either isotype control (10mg/kg/week of mouse anti-IgG antibody) or a mouse anti-IL-1β neutralizing antibody (1mg/kg/week) with similarity to Canakinumab once a week for 8 weeks (**Fig. 5A**). Prior to and following treatment, we performed echocardiography and found a significant improvement in left ventricular ejection fraction (**Fig. 5B**) in *Dsg2*^mut/mut^ mice treated with anti-IL-1β antibody compared to those that received isotype control. Similarly, we observed a decrease in the frequency of premature ventricular contractions and reduced ventricular ectopy in anti-IL-1β antibody treated *Dsg2*^mut/mut^ mice compared to isotype controls (**Fig. 5C**). Upon harvesting hearts from these cohorts, we assessed myocardial fibrosis and found a significant decrease in fibrotic area in anti-IL-1β antibody treated *Dsg2*^mut/mut^ mice relative to isotype-treated counterparts (**Fig. 5D**). Additionally, we performed multiplex cytokine array analysis and found that anti-IL-1β antibody treatment decreased the levels of a number of pro-inflammatory and pro-fibrotic cytokines, including CD14, CXCL2, CXCL9, IFNγ, Osteopontin (OPN), and POSTN (table S4).

**Fig 5.**
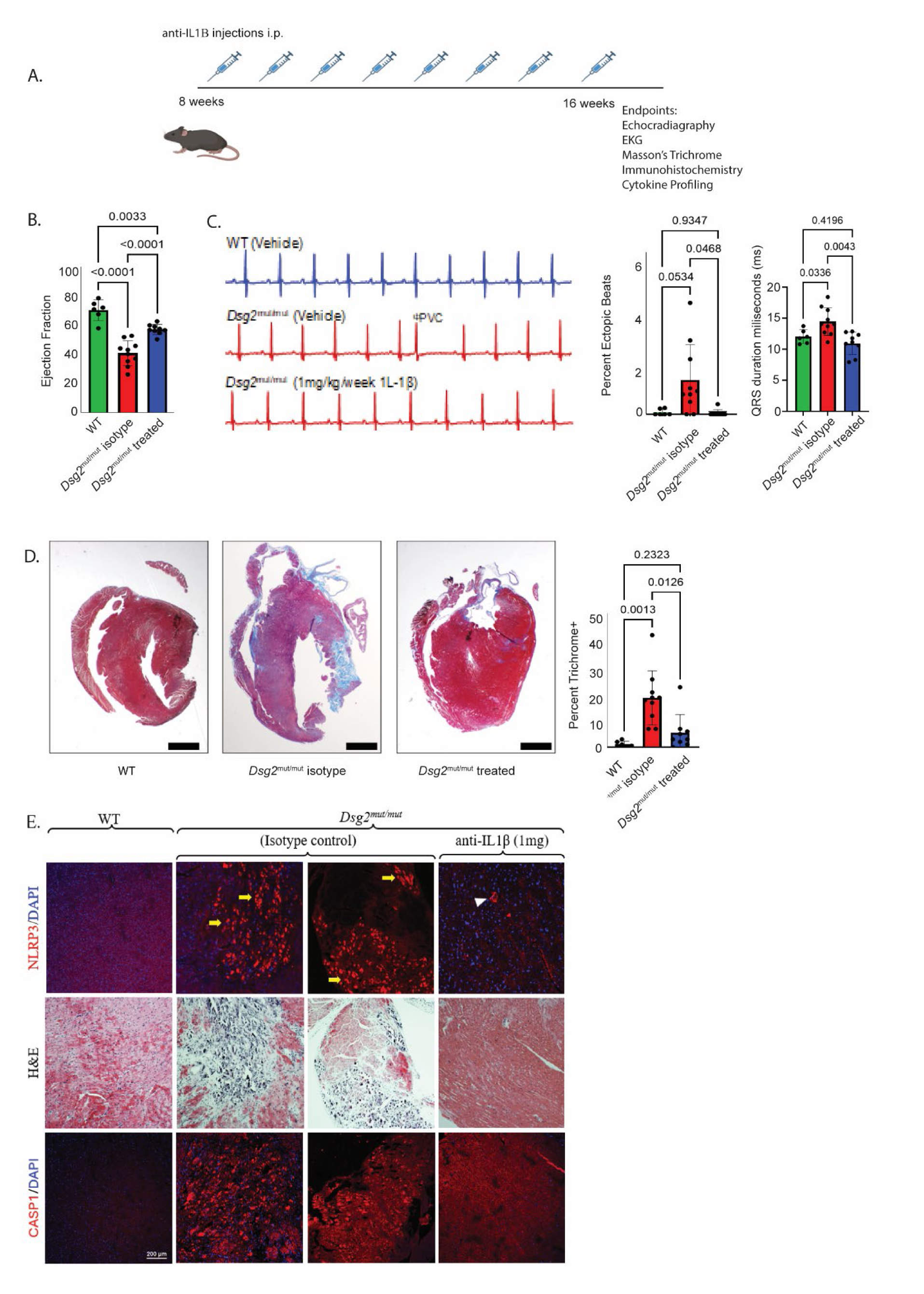
IL1β blockade significantly attenuates disease in *Dsg2^mut/mut^* mice. (**A**) Study design outlining treatment schedule of neutralizing IL1β antibody. (**B**) Measurement of ejection fraction (n=6, 10, and 9 for each group respectively). (**C**) Representative ecgs from each treatment group and quantification of the proportion of ectopic beats and QRS duration (n=6, 10, and 9 for each group respectively). (**D**) Representative Masson’s trichrome images from each treatment group and quantification of fibrosis percentage (n=6, 10, and 9 for each group respectively). Brown-Forsythe and Welch ANOVA testing was used for graphs from (**B**), (**C**), and (**D**). (**E**) Representative immunostained hearts probed for NLRP3, H&E, and CASP1. n≥5 hearts/cohort/stain; yellow arrows, NLRP3 positive staining localized in areas of myocardial lesions; white arrowhead, NLRP3 positive staining localized around vessel lumen.

The NLRP3 inflammasome, a cytosolic complex found in pro-inflammatory immune cells, has been implicated in several cardiovascular diseases and is responsible for the processing and release of IL-1β using active caspase-1 (CASP1) *(26, 34)*. Given our results demonstrating the expansion of *NLRP3*+ macrophages and monocytes in human ACM hearts, we evaluated myocardial tissue for NLRP3 and CASP1 in *Dsg2^mut/mut^*and WT mice. In hearts from isotype-treated *Dsg2^mut/mut^* mice, NLRP3 and CASP1 were exclusively localized in myocardial lesions that additionally showed extensive infiltrating immune cells, a finding not observed in anti-IL-1β antibody treated *Dsg2*^mut/mut^ mice (**Fig. 5E**).

To assess whether a similar benefit could be derived from anti-IL1β antibody treatment at a timepoint in which cardiac dysfunction and biventricular fibrosis are quite evident *(11, 12, 35)*, we treated WT or *Dsg2*^mut/mut^ mice from 16- to 24-weeks of age (fig S6A). At this point of intervention, we found modest improvements in cardiac function in anti-IL-1β antibody treated *Dsg2*^mut/mut^ mice while function deteriorated further in mice that received control antibody (fig S6B). The modest recovery of contractile function in anti-IL1β antibody-treated mice was associated with a substantial reduction in myocardial fibrosis (fig S6C). The more robust improvement in ACM disease at an earlier time point suggests that while anti-IL-1β treatment may have the greatest effect during the early inflammatory stage of disease *(12)*, some benefit can still be achieved at later timepoints. Overall, these findings indicate that therapies targeting inflammation in ACM can provide substantial reductions in disease burden across the natural history of disease.

### Treatment of *Dsg2^mut/mut^* mice with an anti-IL-1β antibody alters the cardiac transcriptional environment

To delineate the transcriptional changes that occur following IL-1β neutralization in *Dsg2^mut/mut^* mice, we performed snRNAseq on 16-week-old WT and *Dsg2^mut/mut^* mice following 8 weeks of either isotype control or anti-IL-1β antibody treatment. We utilized the iCell8cx SMART-seq Pro platform, which allows for the capture of full-length cDNA and a greater number of genes per nuclei (>5,000) compared to other sequencing technologies *(36)*. After performing quality control (fig S7A), we identified six transcriptionally distinct cell types (**Fig. 6A**) marked by major canonically expressed genes (fig S7B). We observed that while the major composition of cell types did not change following anti-IL-1β antibody treatment in *Dsg2^mut/mut^* mice (fig S7C), there was a large number of differentially expressed genes between the anti-IL-1β antibody and isotype-treated *Dsg2^mut/mu^* cohorts in all cell types (**Fig. 6B**). The largest number of differentially expressed genes as well as captured nuclei were in endothelial cells, cardiac myocytes, and fibroblasts. To further investigate changes in the cardiac myocyte gene expression, we performed pathway analysis using the top 25 differentially expressed genes between anti-IL-1β antibody and isotype treated *Dsg2^mut/mut^*hearts. We observed an upregulation in pathways associated with NFĸB-mediated inflammation and cell death in isotype-treated *Dsg2^mut/mut^* mice, while anti-IL-1β antibody-treated cardiac myocytes displayed enrichment in pathways associated with homeostasis and stress response (**Fig. 6C**). These findings are consistent with our previously published observations indicating that NFĸB activity in cardiac myocytes participates in myocardial cell death, cardiac inflammation, arrhythmias, and reduced contractile function *(12)*.

**Fig 6.**
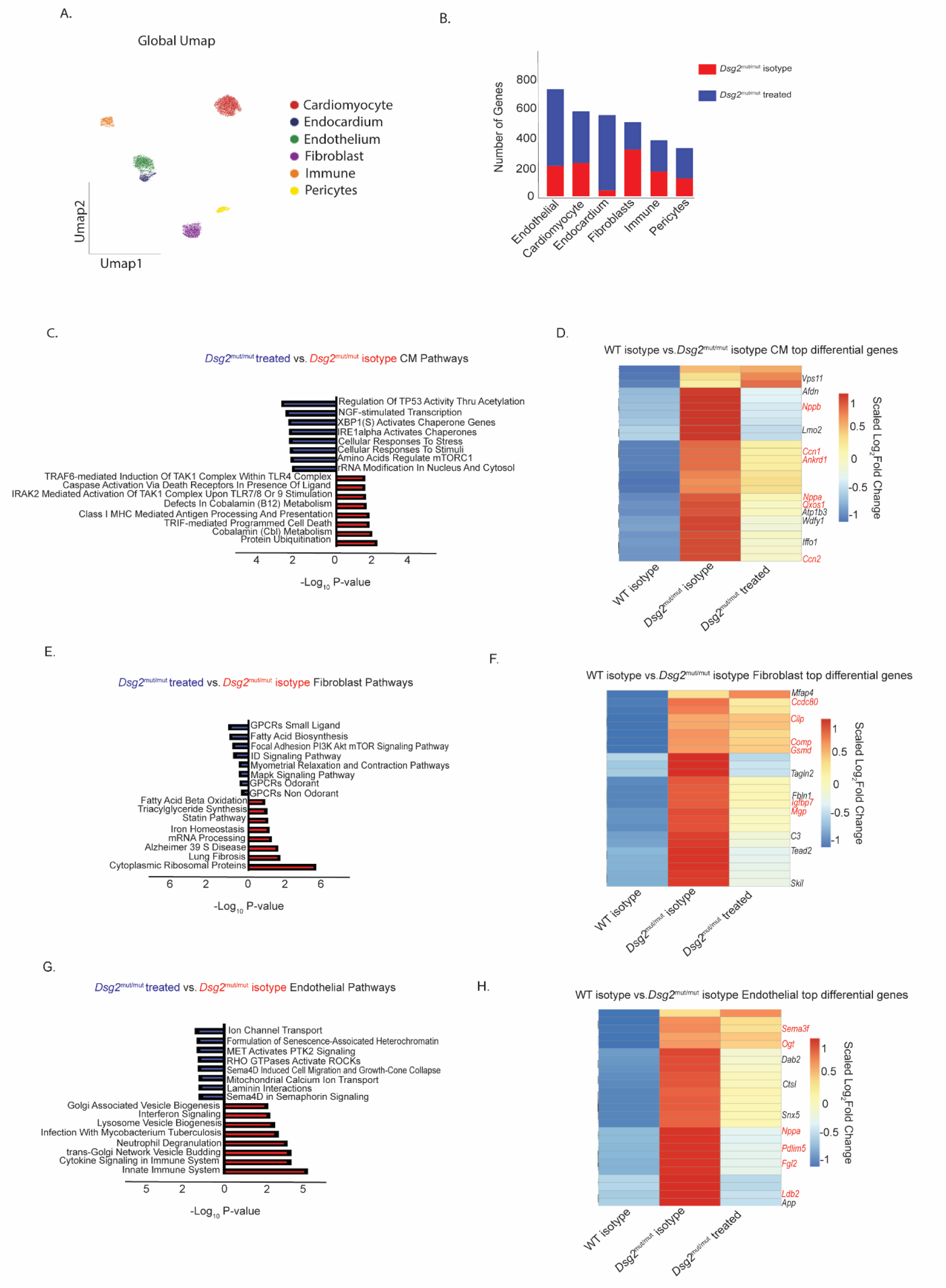
Early IL1β blockade alters the transcriptomic environment in *Dsg2^mut/mut^* mice. (**A**) Global UMAP of cell populations captured in iCell8cx sequencing. (**B**) Total number of differentially expressed genes between *Dsg2^mut/mut^* isotype control and *Dsg2^mut/mut^* anti IL1β treated across different cell types. (**C**) Pathway analysis displaying differentially expressed pathways in cardiac myocytes between *Dsg2^mut/mut^* isotype control and *Dsg2^mut/mut^* anti IL1β treated mice. (**D**) Heatmap displaying top differentially expressed cardiac myocyte genes between WT isotype controls and *Dsg2^mut/mut^* isotype controls and compared to those same genes in *Dsg2^mut/mut^* anti IL1β treated mice. (**E**) Pathway analysis displaying differentially expressed pathways in fibroblasts between *Dsg2^mut/mut^* isotype control and *Dsg2^mut/mut^* anti IL1β treated mice. (**F**) Heatmap displaying top differentially expressed fibroblast genes between WT isotype controls and *Dsg2^mut/mut^* isotype controls and compared to those same genes in *Dsg2^mut/mut^* anti IL1β treated mice. (**G**) Pathway analysis displaying differentially expressed pathways in endothelial cells between *Dsg2^mut/mut^* isotype control and *Dsg2^mut/mut^*anti IL1β treated mice. (**H**) Heatmap displaying top differentially expressed endothelial genes between WT isotype controls and *Dsg2^mut/mut^*isotype controls and compared to those same genes in *Dsg2^mut/mut^*anti IL1β treated mice.

Next, we determined if the broader ACM phenotype had been reversed in cardiac myocytes following anti-IL-1β antibody treatment. To address this question, we determined the expression values of the top 25 differentially expressed genes between WT and *Dsg2^mut/mut^*mouse hearts and assessed whether these genes were impacted by anti-IL-1β antibody treatment in *Dsg2^mut/mut^* hearts (**Fig. 6D**). We observed that a variety of genes upregulated in ACM cardiac myocytes were downregulated following anti-IL-1β antibody treatment, including major heart failure and inflammation associated genes (highlighted in red) *(37–39)*.

We applied the same analysis to the fibroblast and endothelial cell populations and observed an upregulation in pathways associated with fibrosis in isotype-treated *Dsg2^mut/mut^* hearts, while the anti-IL-1β antibody treated fibroblasts displayed enrichment in pathways associated with homeostatic signaling and contraction (**Fig. 6E**). As in cardiac myocytes, many of the major gene signatures upregulated in *Dsg2^mut/mut^* hearts relative to WT were downregulated following treatment, including a number of genes associated with cardiac disease and fibrosis (**Fig. 6F**) *(40–44)*. In endothelial cells, we observed an upregulation in pathways associated with innate immune signaling and inflammation in isotype-treated *Dsg2^mut/mut^* mice, while anti-IL-1β antibody treated endothelial cells displayed enrichment in pathways associated with ion transport and GTPase signaling (**Fig. 6G**). Again, the majority of genes upregulated in *Dsg2^mut/mut^*hearts relative to WT were downregulated following treatment including a number associated with endothelial stress and dysfunction (**Fig. 6H**) *(45–50)*.

### Cardiac myocyte NFĸB nuclear localization and myocardial CCR2/CD68+ infiltrating macrophages is prevented in anti-IL-1β antibody treated *Dsg2^mut/mut^* mice

Given our findings above that demonstrated the areas of myocardial loss harbored macrophages expressing NFĸB-dependent transcripts and a reduction in pathways associated with NFĸB activation in cardiac myocytes, we aimed to determine if cardiac myocyte NFĸB nuclear localization was decreased following treatment. We observed robust immunoperoxidase signal for RelA/p65 in cardiac myocyte nuclei in isotype-treated *Dsg2^mut/mut^* hearts, a finding not observed in WT or anti-IL-1β treated *Dsg2^mut/mut^* myocardium (**Fig. 7A and B**). Additionally, we previously showed that NFκB signaling in cardiac myocytes is liable for mobilizing CCR2+ cells to ACM hearts, where they promote myocardial injury and arrhythmias *(12)*. Accordingly, we assessed the number of infiltrating macrophages via double immunolabeling for CCR2 and CD68 (**Fig. 7A and C**). There was a strong, positive correlation between cells that demonstrated immunoreactivity for both CCR2 and CD68 and those that expressed RelA/p65 in isotype-treated *Dsg2^mut/mut^* hearts, which was absent in anti-IL-1β treated *Dsg2^mut/mut^* hearts (**Fig. 7D**). These findings indicate infiltrating CCR2/CD68+ macrophages in ACM hearts can be blocked via anti-IL-1β treatment.

**Fig 7.**
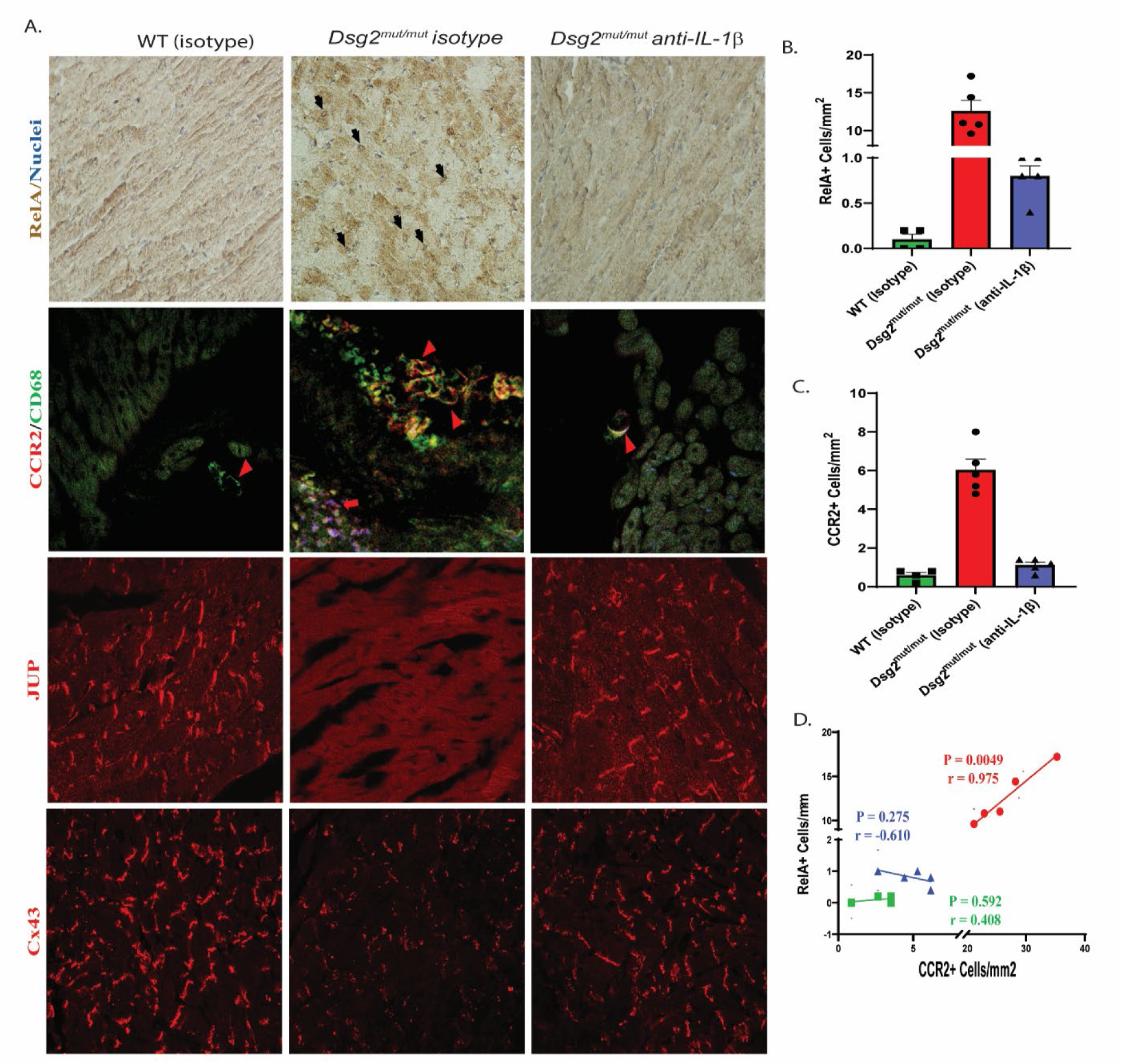
IL1β inhibition prevents NFĸB nuclear localization in cardiac myocytes and infiltrating myocardial CCR2/CD68+ macrophages in *Dsg2^mut/mut^* mice. (**A**) Representative immunostained hearts probed for RelA, CCR2/CD68, JUP, and Cx43. n≥5 hearts/cohort/stain; black arrows, cardiomyocyte RelA positive nuclei; red arrowheads, CCR2/CD68+ macrophages; red arrows: CCR2/CD68+ cardiac myocytes. (**B**) Number of cells per mm^2^ positive for nuclear RelA localization. (**C**) Number of cells per mm^2^ positive for CCR2. (**D**) Pearson’s correlation analysis for cells that showed dual labeling for RelA and CCR2 (P- and r-values inset). For B and C, *P<0.05 for any cohort vs WT+Isotype; and ^†^P<0.05 for any cohort vs *Dsg2^mut/mut^*+Isotype

Loss of junctional plakoglobin (JUP) and connexin-43 (Cx43) at the myocyte-myocyte intercalated disc (ICD) are classical pathological hallmarks of ACM *(51)*. Additionally, recent works reflect that IL-1β released *(52)* from both immune cells and myofibroblasts *(53)* resulted in reduced Cx43 expression and gap junction localization. Therefore, we determined whether the loss of Cx43 at ICD in *Dsg2^mut/mut^* hearts *(11)* could be prevented by anti-IL-1β antibody. Depressed junctional immunoreactive signal for JUP and Cx43 was noted in all *Dsg2^mut/mut^* mice treated with isotype control, compared to WT myocardium (**Fig. 7A**). Abnormal distributions for both proteins were fully corrected in *Dsg2^mut/mut^* mice treated with anti-IL-1β antibody (**Fig. 7A**).

## Discussion

Inflammation and fibrosis are long-recognized features of ACM. Post-mortem explants have revealed significant myocardial fibrosis and inflammatory infiltrates within both the right and left ventricular walls. Moreover, serum levels of inflammatory cytokines, including IL-1β and C-reactive protein (CRP), are elevated in ACM patients *(2, 9, 10, 54)*. Additionally, there has been greater recognition that some individuals diagnosed with acute or recurrent myocarditis carry aggressive ACM mutations, most notably *DSP* variants *(55)*. Recent work employing established mouse models of ACM has provided crucial insights implicating the contributions of inflammation and monocyte recruitment in ACM pathogenesis *(12)*. Despite these advances, the emerging field of cardioimmunology has an incomplete understanding of the immune and stromal landscape of ACM and little is known regarding effector mechanisms that drive myocardial inflammation.

Here we utilized snRNAseq and spatial transcriptomics to define the cellular landscape of human ACM. We uncovered robust expansion of the myeloid and fibroblast populations that localize with ACM lesions. These cell populations harbor the greatest number of differentially expressed genes between donor and ACM hearts. Detailed characterization of fibroblast and myeloid populations expanded in ACM myocardium revealed specific increases in *POSTN* expressing fibroblasts and *NLRP3* expressing inflammatory macrophages. Additionally, we observed a decrease in *F13A1* expressing resident macrophages in ACM hearts compared to donor controls. Similar shifts in fibroblast and macrophage states have been previously reported in myocardial infarction *(23)* and some forms of dilated cardiomyopathy *(19, 24, 25)*. Our results are consistent with and expand upon past ACM sequencing studies *(26)*.

Using spatial transcriptomics, we uncovered the spatial organization of cell types that reside within areas of tissue damage (referred to as ACM lesions). Our data indicated that cardiac myocytes were lost within ACM lesions, which were instead predominantly composed of fibroblasts and macrophages. These lesions were surrounded by niches that contained cardiac myocytes that expressed markers typically observed in heart failure *(19–22)*. ACM lesions resembled an immune-fibrotic niche, which was largely comprised of *POSTN* fibroblasts and inflammatory macrophages. This niche displayed enrichment for inflammatory and fibrotic pathways including NFĸB, TNFα, and TGFβ. Of note, this niche possessed similarities to what is observed in the ischemic and border zones of myocardial infarctions *(23, 28)*, highlighting the potential importance of necroinflammation in ACM.

A key component of inflammatory related processes is crosstalk between immune cells and stromal elements within the tissue microenvironment. Given the proximity of inflammatory macrophages and fibroblasts within ACM lesions, the direct paracrine communication between these cell types may contribute to disease pathogenesis of ACM. To explore a causative relationship between such effector and signaling mechanisms, it is imperative to identify robust models that recapitulate the human disease and are experimentally tractable.

Various *in vitro* and *in vivo* models of ACM have been developed over the years *(56)*, that primarily include models harboring analogous pathogenic gene variants seen in patients with ACM. We leveraged the *Dsg2*^mut/mut^ mouse model, which recapitulates several key pathological and functional aspects of ACM and examined whether analogous macrophage and fibroblast populations exist in this ACM model. Indeed, single cell RNA sequencing of the myeloid and fibroblast populations revealed increases in transcriptionally similar *Postn*+ fibroblasts and inflammatory monocyte and macrophage subsets. These populations expressed many of the genes enriched in human ACM, suggesting that the *Dsg2*^mut/mut^ mouse model could serve as a platform to dissect inflammatory mechanisms that are relevant to human ACM pathogenesis. It is interesting to note that our human sequencing data was derived from patients with *PKP2* or *DSP* variants, which highlights the substantial overlap between gene variants belonging to the cardiac desmosome.

IL-1β is a potent inflammatory cytokine elevated in various forms of cardiac disease and studied in previous clinical trials *(30–32, 57)*. IL-1β plays an important role in adverse remodeling following cardiac injury and promotes myocardial fibrosis via a paracrine communication axis between inflammatory macrophages and fibrotic fibroblasts *(23)*. Although patients with ACM display elevated serum levels of IL-1β *(2, 9, 10)*, its role in ACM pathogenesis is undetermined. Our sequencing data in both human and mouse revealed an increase in NLRP3 and IL-1β expressing macrophages in ACM. Together these findings suggest that IL-1β may be a useful therapeutic target in ACM. Blockade of IL-1β at 8 weeks of age in *Dsg2^mut/mut^* mice using an anti-IL-1β neutralizing antibody, resulted in remarkable improvement in contractile function, decreased fibrosis, and diminished frequency of premature ventricular depolarizations. These findings align with a previously reported study that used an NLRP3 inhibitor in a mouse model of ACM *(26)*.

To determine if targeting IL-1β might be beneficial in advanced disease, we also blocked IL-1β in 16-week-old *Dsg2*^mut/mut^ mice. We observed modest improvements in ejection fraction and a substantial decrease in cardiac fibrosis. These findings suggest benefits across the natural history of ACM with greater efficacy at earlier timepoints, which are hypothesized to resemble periods of greater inflammation *(12)*. In mouse models, monocytes appear to peak at 4-6 weeks of age *(12)*. Nevertheless, there is a large degree of heterogeneity in the progression of ACM, especially in humans *(1)*. Imaging cardiac inflammation using novel PET tracers may be a useful strategy for identifying which patients would benefit most from anti-inflammatory intervention *(58, 59)*.

To uncover potential transcriptional changes in the cardiac environment following treatment with anti-IL-1β antibody, we performed snRNAseq on 16-week-old WT and *Dsg2^mut/mut^* treated hearts using the iCell8cx SMART-seq Pro platform to leverage its ability to obtain more genes per nuclei. Analysis of the sequencing results revealed a plethora of differentially expressed genes. Within cardiac myocytes, we observed substantial reductions in the ACM cardiac gene signature in anti-IL-1β antibody treated ACM mutants. Using pathway analysis, we observed downregulation of NFĸB-induced inflammation and cell death associated pathways in following IL-1β antibody treated ACM mutants. We have previously demonstrated that NFĸB-dependent cell death in cardiac myocytes is a major determinant regulating the recruitment of inflammatory monocytes in ACM hearts via transcriptional upregulation of potent chemotactic molecules *(12)*. IL-1β blockade has been shown to reduce cardiac fibrosis *(60)*, intestinal cell death in a model of small intestine enteropathy *(61)*, and beta islet cell death in a rat model of type I diabetes *(62)*. The reduction of cardiac myocyte cell death, fibrosis, and inflammation highlights a probable mechanism of action by which IL-1β blockade attenuates ACM pathogenesis.

We recognize that our study is not without limitations. The human myocardial samples studied were obtained from patients at time of heart transplantation and thus, these hearts were collected during advanced stages of disease progression. As a result, we were unable to distinguish the cellular and transcriptional landscape in ACM hearts during different stages of disease (i.e., “Concealed” vs “Hot” Phases) at time of sequencing. Given the heterogenous disease course and variable age of diagnosis, this may have an impact on the cellular populations and transcriptional states we detected. Additionally, while we sequenced hearts harboring two of the most prevalent genes that give rise to ACM, we recognize that additional pathogenic variants in several other genes are associated with an ACM-like phenotype *(5)*. Although our data suggests that there is some degree of conservation across desmosomal genes, it is possible that pathogenic variants in structural or Z-disc genes may display distinct features. We also recognize that our myocardial samples come entirely from the left ventricle of ACM patients. Adipocyte dysplasia, observed primarily in the right ventricle *(63)*, is a major clinical finding in ACM *(1, 2, 4)*, therefore a similar analysis focused on right ventricular myocardial samples would be fruitful in understanding adipocyte contribution to ACM pathogenesis. Lastly, while we focus on the role of IL-1β, it is likely that other inflammatory signaling pathways contribute to ACM.

In summary, we observe a conserved expansion of inflammatory macrophages and fibrotic fibroblasts in ACM in both mice and humans. These cell types are spatially localized to ACM marked areas of tissue damage and fibrosis (i.e., lesions). We demonstrate that IL-1β produced by pro-inflammatory macrophages participates in ACM pathogenesis by driving inflammation, fibrosis, contractile dysfunction, and an arrhythmogenic substrate. Our findings highlight the utility of anti-inflammatory therapies for ACM, which may serve as a new avenue to treat this devastating disease.

## Materials and Methods

### Study Design

The objective of this study was to elucidate the cellular and transcriptional environment of ACM, and subsequently to better understand the role of inflammatory mediators (specifically IL-1β) on ACM disease progression. To assess the former, we performed snRNAseq and spatial transcriptomics using the 10x Genomics 5’v2 and Visium platforms, respectively. All human myocardial samples were previously frozen and obtained from the Tissue Cardiovascular Biobank and Repository at Washington University. For snRNAseq, we utilized all donor and ACM samples that met quality control cutoffs following library construction. For spatial transcriptomics, we selected tissue samples that met RNA quality control standards based on DV200 measurements prior to sectioning and sequencing *(16, 17)*. For animal studies, we used the previously established *Dsg2^mut/mut^* mouse model *(11)*. To assess the presence of analogous myeloid and fibroblast populations in mice, we performed snRNAseq on sorted myeloid cells and fibroblasts from WT and *Dsg2^mut/mut^*mice. The effects of IL-1β blockade on ACM disease progression in these mice was assessed via echocardiography, electrocardiography, histology, cytokine proteome arrays, and snRNAseq. Investigators were blinded to both genotype and drug treatment cohorts throughout experiments and analyses. Sample size (*n* = 5 to 9) was predetermined based on our prior publications, *(11, 12, 29)* which demonstrated n≥4 mice was more than sufficient to show substantial cardiovascular and pathological abnormalities between cohorts (i.e., WT vs. *Dsg2*^mut/mut^). Utilizing n≥4 mice/genotype/cohort/parameter, we were able to achieve an 80–99% Power Analyses and receive a statistical probability of P<0.05. No data was excluded from any analyses. Further details on specific methods are described in individual sections below.

### Ethical approval for human samples

This study is compliant with all relevant ethical regulations and approved by Washington University School of Medicine Institutional Review Board (IRB no. 201104172). Each patient provided informed consent prior to tissue collection and no compensation was provided for study participation. All patients have been deidentified.

### Human sample inclusion criteria

Donor myocardial samples were obtained from patients classified with stable ejection fractions, no known history of cardiac disease, and who experienced a non-cardiac related reason for transplant or death were provided by Mid America Transplant Service. ACM myocardial samples were derived from patients undergoing heart transplant. Specific ACM pathogenic variants were determined for each sample using the ClinVar database. Only samples that had a single gene variant for ACM were included in this study.

### Animals

All experiments conformed to the Guide for the Care and Use of Laboratory Animals from the National Institute of Health (NIH publication no. 85–23, revised 1996). Animal study protocols were approved by the Florida State University (Protocol Code: 202000052; Date of Approval: 02/10/2021) and Washington University in St. Louis (Protocol Code: D1600245, Date of Approval: 12/02/2020) Animal Care and Use Committee. Mice were housed in temperature-controlled rooms (20–22°C) and humidity (40–60%) with a 12hr light/dark cycle and provided *ad libitum* access to standard rodent chow and water. Age-matched C57BL/6 mice served as WT controls. The generation of *Dsg2^mut/mut^* has been previously described *(11)*.

### Human myocardial tissue processing

Several 7-10 mm^3^ sized pieces were collected from the left ventricle of explanted hearts. Some were flash frozen in liquid nitrogen to be used for downstream RNA sequencing studies, while others were fixed in 4% paraformaldehyde and then embedded in paraffin for histology and spatial transcriptomics.

### Human 10x single nuclei processing

Frozen human myocardial samples were prepared for single nuclei RNA sequencing (snRNAseq) as described previously *(19)*. In brief, tissues were minced with a razor blade and transferred to a Dounce homogenizer containing lysis buffer (10x Genomics). Samples were homogenized gently and then incubated on ice for 15 min. Following filtration and washing steps, samples were stained with 1ul of DRAQ5 (Thermo Scientific Cat No: 62251) and DRAQ5+ nuclei were sorted and counted on a hemocytometer (Thermofisher). Nuclei were subsequently processed using the Chromium Single Cell 5’ Reagent V2 kit from 10x Genomics (PN-1000263). A total of 10,000 nuclei per sample were loaded onto the chip for GEM generation. Library preparation was performed according to protocol and sequencing was performed on a NovaSeq 6000 platform (Illumina).

### Human 10x Visium slide preparation

Human myocardial samples were paraffin-embedded as described above. To determine tissue RNA quality, DV200 values for each sample were determined using the RNeasy FFPE kit (Qiagen Cat No:73504). Samples with a DV200 value greater than 40 were selected for downstream processing. 10 µm sections were placed on spatial gene expression slides (Visium, 10x Genomics, PN-1000187). Samples were processed following the Visium User Guide (CG000239). Brightfield histology images were taken on a Zeiss Axioscan Z1 (Carl Zeiss AG). Libraries were generated according to the Visium user guide and sequenced on a NovaSeq 6000 (Illumina).

### Mouse 10x Single Cell preparation

Samples were prepared as described previously *(12)*. In brief, freshly isolated hearts from 6-week-old WT and *Dsg2^mut/mut^* mice (n=3 per group) were minced on ice with a razor blade, transferred to a 15ml conical tube containing 3mL DMEM (Gibco Cat No: 11965-084) with 170µL collagenase IV (Sigma Cat No: C5138-1G) (250U/mL final concentration), 35µL DNAse1(Sigma Cat No: D4527-40KU) (60U/mL), and 75µL hyaluronidase (Sigma Cat No: H3506) (60U/mL), and incubated at 37°C for 40 min with gentle agitation. The digestion reaction was then quenched and filtered. Samples were then centrifuged at 4°C, for 5 min at 1200 rpm and the supernatant was discarded. Pellets were resuspended in 1mL ACK Lysis buffer (Gibco, Cat. No. A10492-01) and incubated at room temperature for 5 min, then quenched with DMEM and centrifuged as above. Supernatant was discarded and the pellets were resuspended in 1mL of FACS buffer and divided evenly by volume before 500µl of respective antibody staining cocktail (for fibroblasts and myeloid) was added (data supplement). Samples were stained for 30 min at 4 degrees in the dark. Samples were washed with FACS buffer and centrifuged as above and resuspended in 300ul of FACS buffer. Samples were sorted on a BD FACS Melody and subsequently counted on a hemocytometer. Fibroblasts were identified as PDGRFA+PDPN+CD45-CD31-. Myeloid cells were identified as CD45+CD11b+Ly6G-. Collected cells were processed using the Single Cell 3’ v3.1 kit (10x Genomics PN: 1000268). 10,000 cells from each sample were loaded onto the chip for GEM generation. Library preparation was performed according to protocol and sequencing was performed on a NovaSeq 6000 platform (Illumina).

### 10x single nuclei RNA seq analysis

Alignment, quality control, and filtering were performed as previously described *(19)*. In brief nuclei were aligned to the human GRCh38 reference using CellRanger v6.1. Subsequent quality control, normalization, dimensional reduction, and clustering were performed in Seurat v4.0. Following normalization, quality control was performed and cells passing the following criteria were kept for downstream processing: 500 < nFeature_RNA < 4000 and 1,000 < nCount_RNA < 10,000 and percentage mitochondrial reads < 5%. To remove doublets, Scrublet was run with a cutoff score of >0.25 to identify doublets. Following doublet removal, raw RNA counts were normalized and scaled using SCTransform. PCs were then calculated, and an elbow plot was generated to select the cutoff for significant PCs to use for downstream analysis. UMAP dimensional reduction was then computed using the selected significant PCs. Unsupervised clustering was then performed using the FindNeighbors and FindClusters function, again using the selected significant PC level as above, calculating clustering at a range of resolutions between 0.1–0.8 at intervals of 0.1. Differential gene expression was performed using the FindAllMarkers command and a Wilcoxon Rank Sum test with a logFC cutoff of 0.25 and a min.pct cut-off of 0.1. Clusters were annotated using canonical gene and protein markers. To cluster cell types into distinct cell states, the cell type of interest was subsetted, renormalized, computed PCAs, computed UMAPs, and clustered data at a range of resolutions. DE analysis was then used to identify marker genes for each cell state. Using the top marker genes, gene set z-scores were calculated and plotted in UMAP space. Pseudobulk differential gene expression was performed as pervious described *(23)*.

### Visium Spatial Transcriptomics Analysis

Space ranger was used to align counts, and the following matrices were processed using Seurat v4, stlearn, and tangram. First, tangram was used in conjunction with the snRNA-seq atlas for voxel deconvolution using baseline parameters. Cell type deconvolution scores were overlaid on the spatial histological images to infer spatial localization. Within the Seuratv4 and stlearn pipeline, the data was normalized and clustered with dimensional reduction to identify spatial niches which were characterized using differential gene expression analysis. Spatial correlation analysis was used to identify cell types which co-localize in space – briefly, the tangram deconvolution score matrix was used and a correlation matrix constructed for downstream visualization (for the purposes of the visualization the diagonal was set to 0). For building pie charts in space, we used the tangram deconvolution scores and stlearn to overlap compositional pie charts in space. Progeny was used for pathway analysis across spatial niches.

### 10x single cell RNA seq analysis

Alignment, quality control, and filtering were performed as previously described *(12, 23)*. Cells were aligned to the mouse GRCh38 using CellRanger v6.1. Subsequent filtering and quality control was performed as above using the following filters for fibroblasts and myeloid cells respectively: 500 < nFeature_RNA < 6000 and 1,000 < nCount_RNA < 25,000 and percentage mitochondrial reads < 5%, 500 < nFeature_RNA < 6000 and 1,000 < nCount_RNA < 25,000 and percentage mitochondrial reads < 10%. Normalization and clustering were performed as above.

### Pathway analysis

Pathway analysis was performed as previously described using EnrichR (http://maayanlab.cloud/Enrichr/) *(12, 23)*.

### Immunofluorescence staining

Human myocardial samples were fixed overnight in 4% paraformaldehyde and then embedded in paraffin blocks. 10 um sections were cut and placed onto charged slides. For immunofluorescence staining we followed the Opal IHC assay guide (Akoya Biosciences) and used the following primary antibodies: anti-CD68 (BioRad clone KP1), anti-CCR2 (Abam clone 7A7), and anti-fibroblast activation protein (Abcam clone EPR20021). Images were captured on a Zeiss Axioscan 7 and analyzed in Zen Blue.

### Immunoperoxidase staining

Formalin-fixed, paraffin-embedded mouse hearts were analyzed by immunoperoxidase staining using a primary antibody against RelA/p65. Sections (5μm thick) were deparaffinized, dehydrated, rehydrated, and exposed to 3% hydrogen peroxide solution for 10 min to block endogenous peroxidase activity. Sections were first incubated with 5% donkey serum (Stratech Scientific, Cat. No. 017-000-121) and 5% BSA (Merck Life Science, Cat. No. A2153) in 1X TBS blocking solution for 1hr then overnight at 4°C with rabbit anti-RelA polyclonal antibody (LSBiosciences Cat. No. LS-B653; at 1:100). The following day, sections were incubated with horseradish peroxidase donkey anti-rabbit secondary antibody (ThermoFisher Scientific, Cat. No. A16038, at 1:400) for 1hr at room temperature. Peroxidase-conjugated antibodies were detected by the 3,3’-diaminobenzidine (DAB) substrate kit (AbCam, Cat. No. ab64238). Slides were then counterstained with Mayer’s hematoxylin (CELLAVISION, Cat. No. 361075) and bright field images were taken with a Nikon Eclipse 80i microscope and recorded with Nikon DS-Fi1 camera. The number of cardiac myocytes showing nuclear signal for RelA were counted in 5 regions of interest (ROIs) and expressed as the number of cells/mm^2^.

### Single-labeling immunofluorescence staining

Mouse myocardial sections were analyzed by immunofluorescence staining using primary antibodies against junctional plakoglobin (JUP) and connexin-43 (Cx43). Sections (5μm thick) were deparaffinized, dehydrated, rehydrated, and boiled in citrate buffer (pH 6) for 11 min. Slides were incubated with 3% goat blocking solution (Stratech Scientific, Cat. No. 005-000-121), containing 1% BSA (Merck Life Science, Cat. No. A2153) and 0.15% Triton-X (ThermoFisher Cat. No. A16046.0F) in 1X PBS for 1hr and then rabbit anti-JUP monoclonal antibody (AbCam, Cat. No. Ab184919, at 1:100) or rabbit anti-Cx43 polyclonal antibody (Sigma Aldrich, Cat. No. C6219, at 1:200) overnight at 4°C. The following day, sections were incubated with SPECIES anti-rabbit Cy3-labeled secondary antibody and mounted with ProLong Gold. Images were obtained using a Nikon A1R confocal microscope.

### Double-labeling immunofluorescence staining

Mouse myocardial sections were analyzed by double-labeling immunofluorescence staining using primary antibodies against CCR2 and CD68. Briefly, sections (5μm thick) were incubated at 60° (20mins), deparaffinized, dehydrated, rehydrated and boiled in citrate buffer (pH 6) for 11min. Slides were then washed in 1X PBS (3x, 5mins/wash), incubated with 5% donkey serum (Stratech Scientific, Cat. No. 017-000-121), 1% BSA (Merck Life Science, Cat. No. A2153) and 0.15% Triton-X (ThermoFisher, Cat. No. A16046.0F) in 1X PBS for 1hr and then probed with rat anti-CD68 antibody (ThermoFisher, Cat. No. 14-0681-82, at 1:200) and rabbit anti-CCR2 antibody (ThermoFisher, Cat. No. BS-23026R, at 1:400) overnight at 4°. The following day, sections were incubated with SPECIES anti-rat FITC-conjugated and SPECIES anti-rabbit Cy5-conjugated secondary antibodies and mounted with ProLong Gold (ThermoFisher, Cat. No. P10144). Images were obtained using a Nikon A1R confocal microscope. The number of macrophages showing strong immunofluorescent signal for CCR2 were counted in 5 ROIs and expressed as the number of cells/mm^2^.

### IL1β treatment

Mouse anti-IL1β and isotype control (mouse anti-IgG) were generously provided by Novartis. WT and *Dsg2^mut/mut^* mice were randomly assigned to either the isotype control or anti-IL1β treatment groups at 8 weeks of age (early intervention) or 16 weeks of age (late intervention). Mice in both studies were treated for 8 weeks with either 1mg/kg/week of mouse anti-IL1β or 10mg/kg/week of isotype control via hindlimb intramuscular injection.

### Animal Echocardiography

Cardiac function was assessed prior to and at treatment endpoint using the Vevo F2 Imaging Platform (Fujifilm Visualsonics, Washington). The Vevo F2 imager was utilized to obtain both short- and long-axis images at the level of the papillary muscles (sweep speed of 200 mm/s), as previously described *(11, 35)*. Images were analyzed using the American Society of Echocardiography guidelines for animals *(64)*. Three to five images were obtained for each mouse/timepoint, then averaged to assess %LVEF and wall/chamber dimensions.

### Animal ECGs

Mice were anesthetized via nose cone anesthesia (1.5 - 2% isoflurane vaporized in 100% O_2_) and ECG electrodes were placed between the right and left front paws while the mouse was in a supine position to obtain Lead I ECG recordings, as previously described *(35)*. iWorx 8 lead channel (iWorx Bio-8, New Hampshire) with ECG Analysis Add-on Software was utilized to analyze Signal-averaged ECGs (SAECGs) from 10-minute recordings. ECG Analysis Add-on Software afforded us to measure the following parameters: wave durations or intervals (via SAECGs) and wave amplitudes (via SAECGs). Percent premature ectopic contractions (%PVCs) were analyzed via totaling the number PVCs throughout the 10-minute recording then divided by the number of total beats, times 100. Following terminal ECG recordings (i.e., at 16 or 24 weeks of age), mice were euthanized and hearts excised for downstream pathological, protein/mRNA and sequencing analyses.

### Masson’s Trichrome Immunostaining

Hearts were formalin-fixed, paraffin-embedded (FFPE) and blocks cut at 5µm (2-3 cuts per slide) and stained with Masson’s Trichrome stain (Sigma; Cat. No. HT15-1KT) following manufacturer’s protocol. Myocardial sections were traced using ImageJ to achieve total area. Then blue fibrotic sections were traced to record total fibrosis. Percent myocardial fibrosis was determined by the sum of all fibrotic areas (within one slice) divided by total myocardial area using ImageJ version 1.53e software. Each slice was then averaged to obtain percent myocardial fibrosis for one mouse.

### Mouse single nuclei preparation for iCellcx8 sequencing

Nuclei were isolated from frozen mouse heart samples from the early intervention IL1β treatment experiment using the Chromium Nuclei Isolation kit from 10x Genomics (PN: 100047), stained with DRAQ5. Following staining, DRAQ5+ nuclei were sorted and counted on a hemocytometer. Subsequently we utilized and followed the SMART-Seq Pro Application kit (Takara Biosciences) to prepare samples for dispense by the iCell8cx system and subsequent reverse transcription, cDNA amplification, and library generation. Libraries were sequenced on one full lane of NovaSeq X Plus.

### iCellcx8 library alignment

iCellcx8 sequencing data was aligned to the mouse reference genome via Cogent AP (Takara Biosciences). Following alignment gene matrix files were generated for each library. Gene matrices were converted into an acceptable format for Seuratv4 on R. Subsequent analysis was performed as above for single nuclei samples.

### Statistical analysis

Data is presented as mean ± SEM (n-values inset within figure legend) for all in vivo studies. Brown-Forsythe and Welch ANOVA was used for comparisons between multiple groups with unequal variances. P < 0.05 was considered significant. All results were repeated at least twice under the same or similar experimental conditions. All statistical analyses were done using GraphPad Prism (v10) software.

## List of Supplementary Materials

figs S1 to S7

tables S1 to S4

Data supplement file

## Acknowledgments

We thank the Genome Technology Access Center at the McDonnell Genome Institute at Washington University School of Medicine for their help with sequencing and genomic analysis. Figs 1a, 5a, and S5a were created using BioRender. We thank Novartis for generously providing the anti-IL-1β antibody and isotype control antibody.

## Funding

National Institutes of Health 5T32AI007163-44 (VRP)

American Heart Association Predoctoral Fellowship 826325 (JMA)

American Heart Association Career Development Award 19CDA34760185 (SPC)

Florida State University Institute of Pediatric Rare Diseases (SPC)

National Institutes of Health grant R01-HL148348 (JES)

Washington University in St. Louis Rheumatic Diseases Research Resource-Based Center grant NIH P30AR073752 (KJL)

National Institutes of Health grant R35-HL161185 (KJL)

Leducq Foundation Network grant #20CVD02 (KJL)

Burroughs Wellcome Fund grant 1014782 (KJL)

Children’s Discovery Institute of Washington University and St. Louis Children’s Hospital grant CH-II-2015-462, Ch-II-2017-628, PM-LI-2019-829 (KJL)

Foundation of Barnes-Jewish Hospital grant 8038-88 (KJL)

## Author Contributions

Conceptualization: KJL, JES, SPC

Experiments: VRP, JMA, PM, AV, AP, ME, WF, AA, CBB, SPC, JJ, CLT

Data analysis: VRP, JMA, SPC, AA, HS

Writing original draft: VRP, JMA, SPC

Writing review and editing: VRP, JMA, SPC, JES, AA, KJL

## Data availability

Raw and processed sequencing files will be uploaded to the Gene Expression Omnibus. Code is available upon request to the authors.

