## Supplemental materials for "Interleukin-1β Drives Disease Progression in Arrhythmogenic Cardiomyopathy"

### List of Supplemental Materials

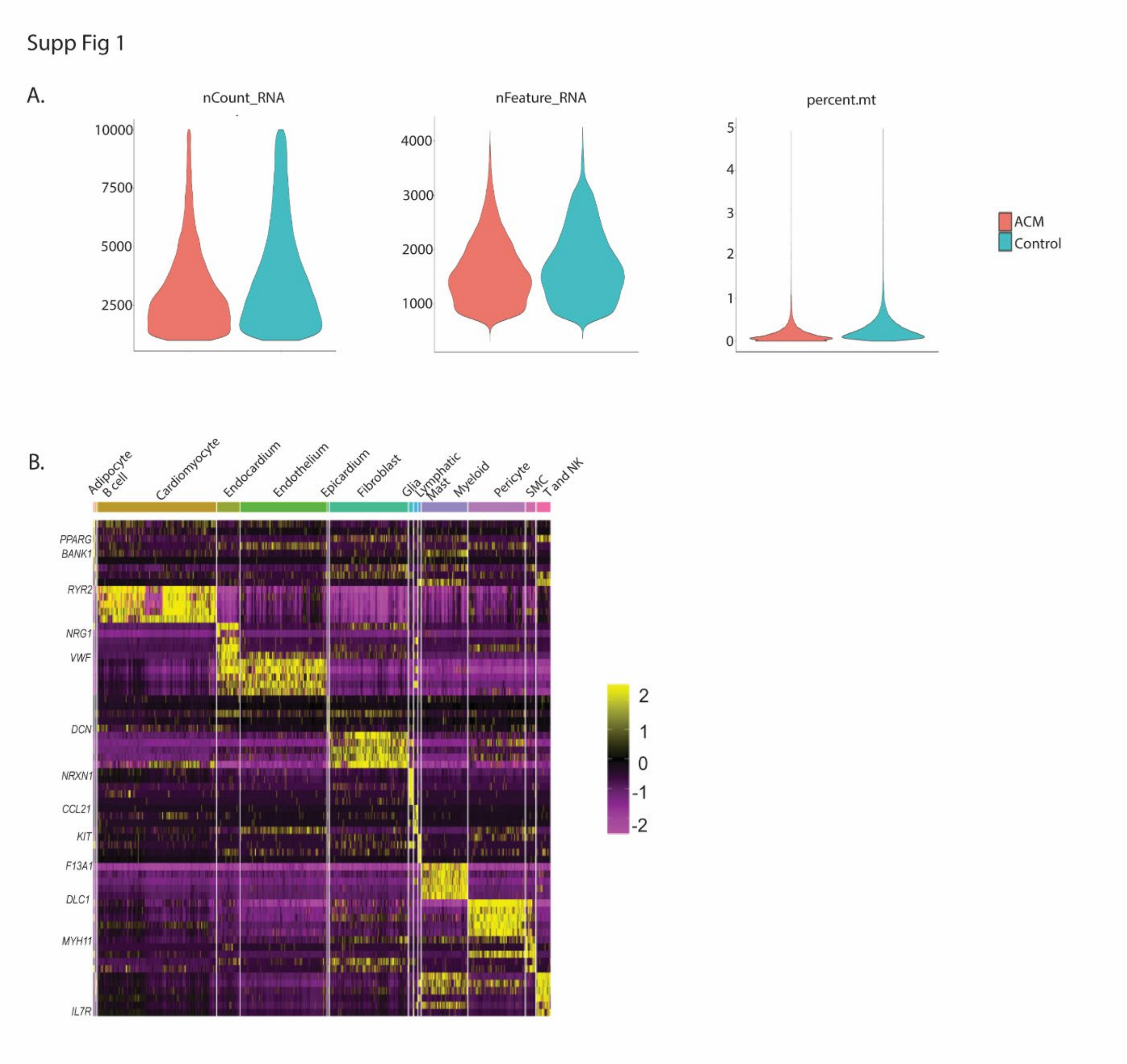


**fig S1 QC metrics for human myocardial samples.**

(**A**) Major QC cutoffs for single nuclei RNA sequencing data. RNA count: 1000<n<10000, Feature count: 500<n<4000, Mitochondrial percentage: n<5. (**B**) Heatmap displaying major canonical gene expression across different cell types.

Sup Fig 2

Donor 1

Donor 2

ACM 1

ACM 2

ACM 3

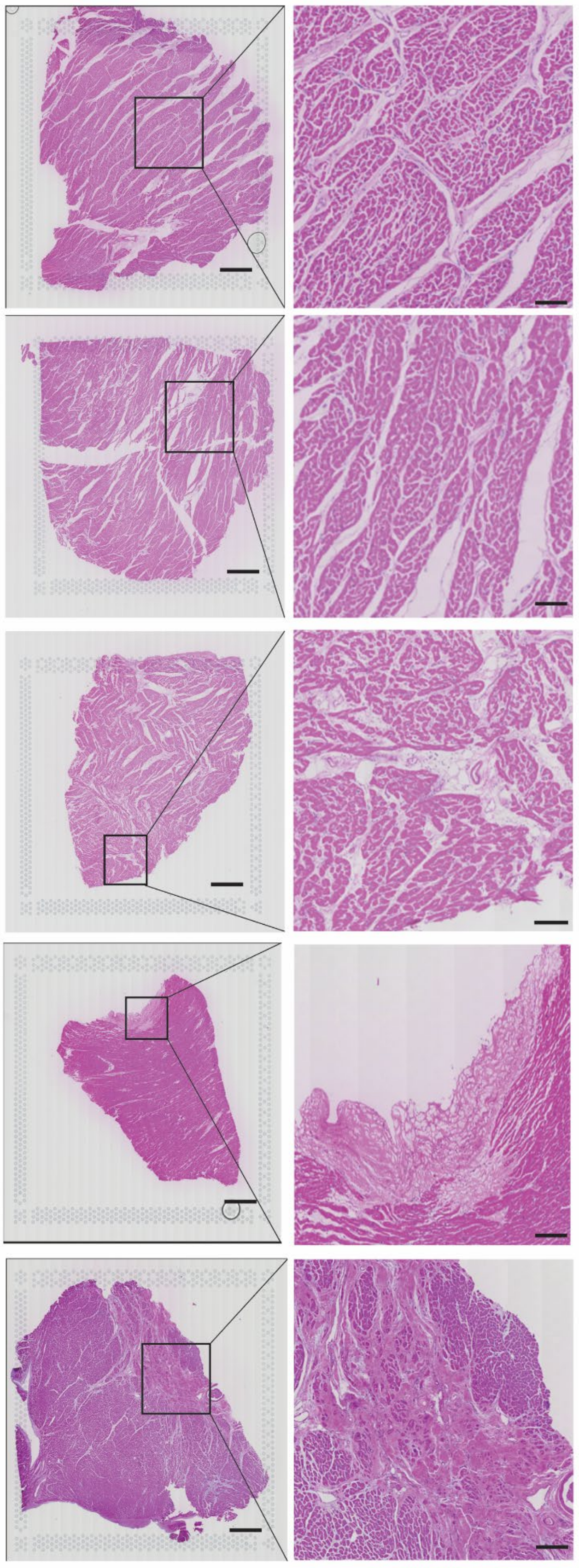

**Fig S2 H&E images used for Visium spatial transcriptomics**

H&E images for each sample used for spatial transcriptomics. Images to the left were captured on an Axioscan Z7. Images to the right are zoomed insets of areas of healthy myocardium (in donor samples) or areas of ACM lesions (in ACM samples).

Supp Fig 3

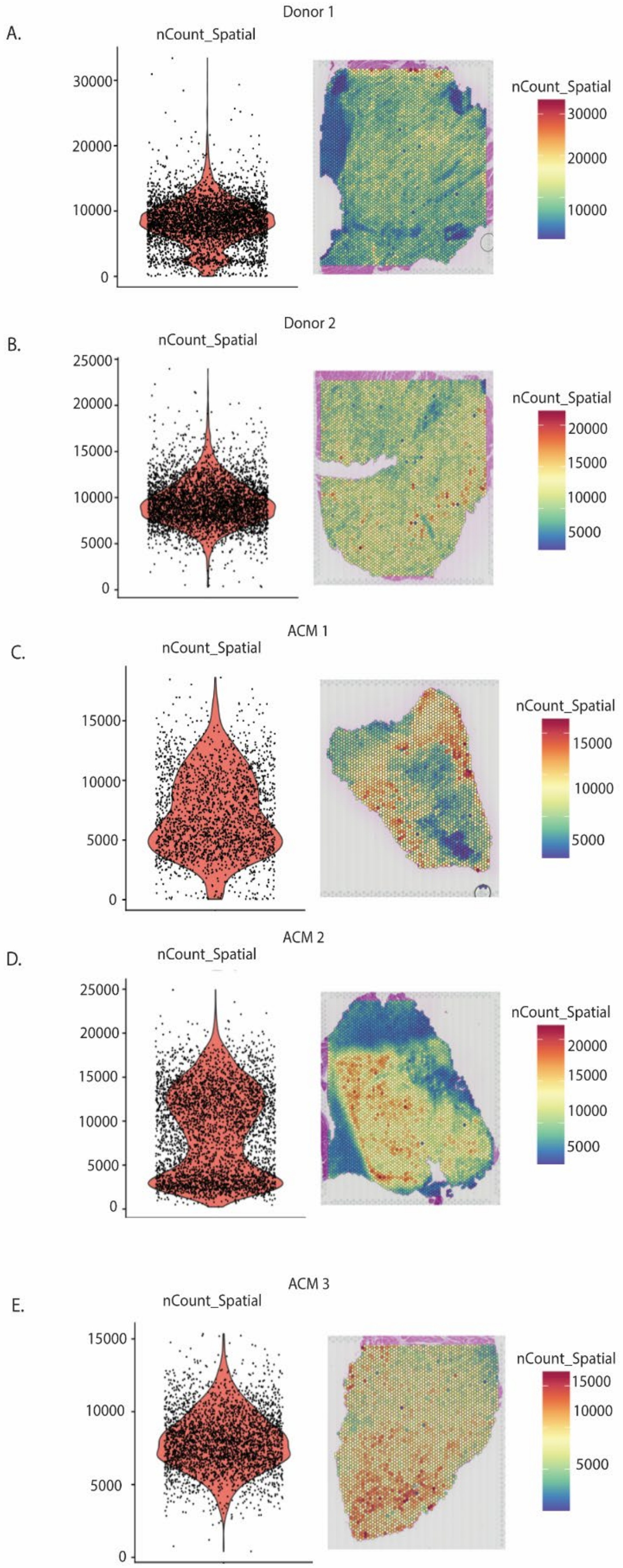

A.
Donor 1
nCount_Spatial
30000
20000
10000
0
nCount_Spatial
30000
20000
10000
B.
Donor 2
nCount_Spatial
25000
20000
15000
10000
5000
0
nCount_Spatial
20000
15000
10000
5000
C.
ACM 1
nCount_Spatial
15000
10000
5000
0
nCount_Spatial
15000
10000
5000
D.
ACM 2
nCount_Spatial
25000
20000
15000
10000
5000
0
nCount_Spatial
20000
15000
10000
5000
E.
ACM 3
nCount_Spatial
15000
10000
5000
0
nCount_Spatial
15000
10000
5000

**Fig S3 QC metrics for spatial transcriptomics samples.**

Spatial UMI counts for each sample: (**A**) Donor 1, (**B**) Donor 2, (**C**) ACM 1, (**D**) ACM 2, (**E**) ACM 3.

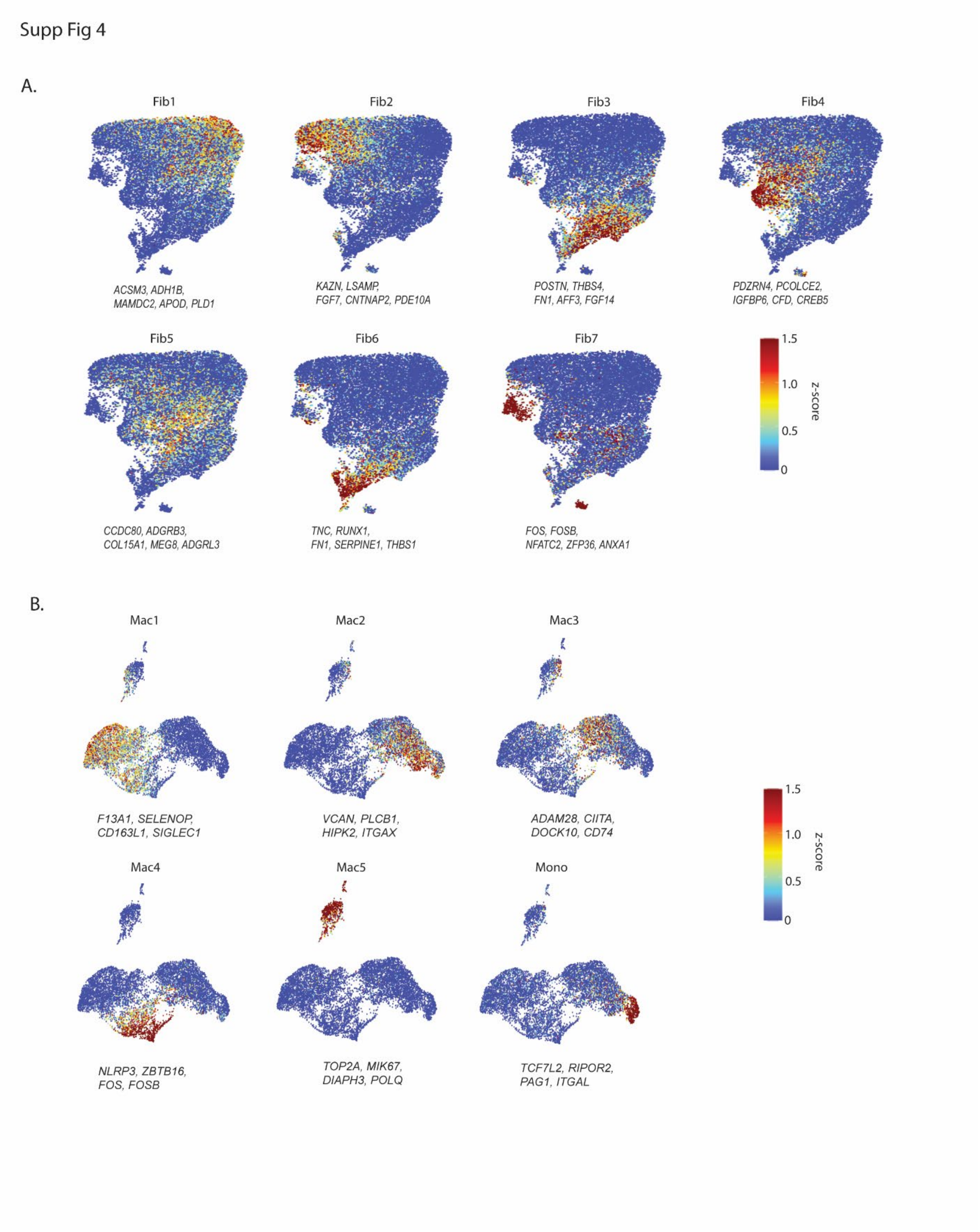


**Fig S4 Sub population z-scores for human fibroblasts and myeloid cells.**

(**A**) Fibroblast sub populations identified by major gene expression. (**B**) Myeloid sub populations identified by major gene expression.

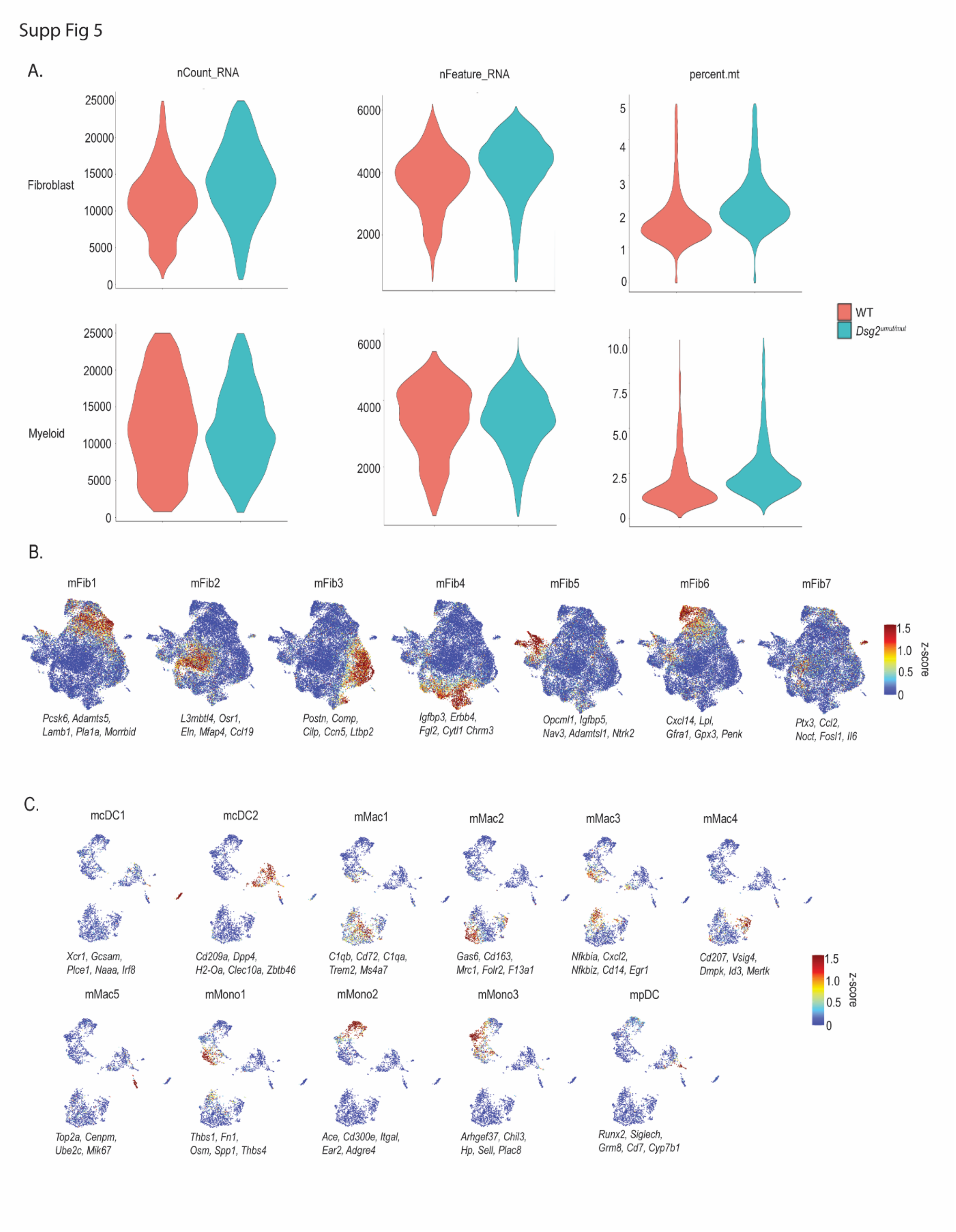


**Fig S5 Mouse single cell RNA sequencing QC metrics and major population z-scores.**

(**A**) Major QC cutoffs for single cell RNA sequencing data. RNA count: 1000<n<25000, Feature count: 500<n<6000, Mitochondrial percentage: n<5 for fibroblasts and n<10 for myeloid. (**B**) Mouse fibroblast sub populations identified by major gene expression. (**C**) Mouse myeloid sub populations identified by major gene expression.

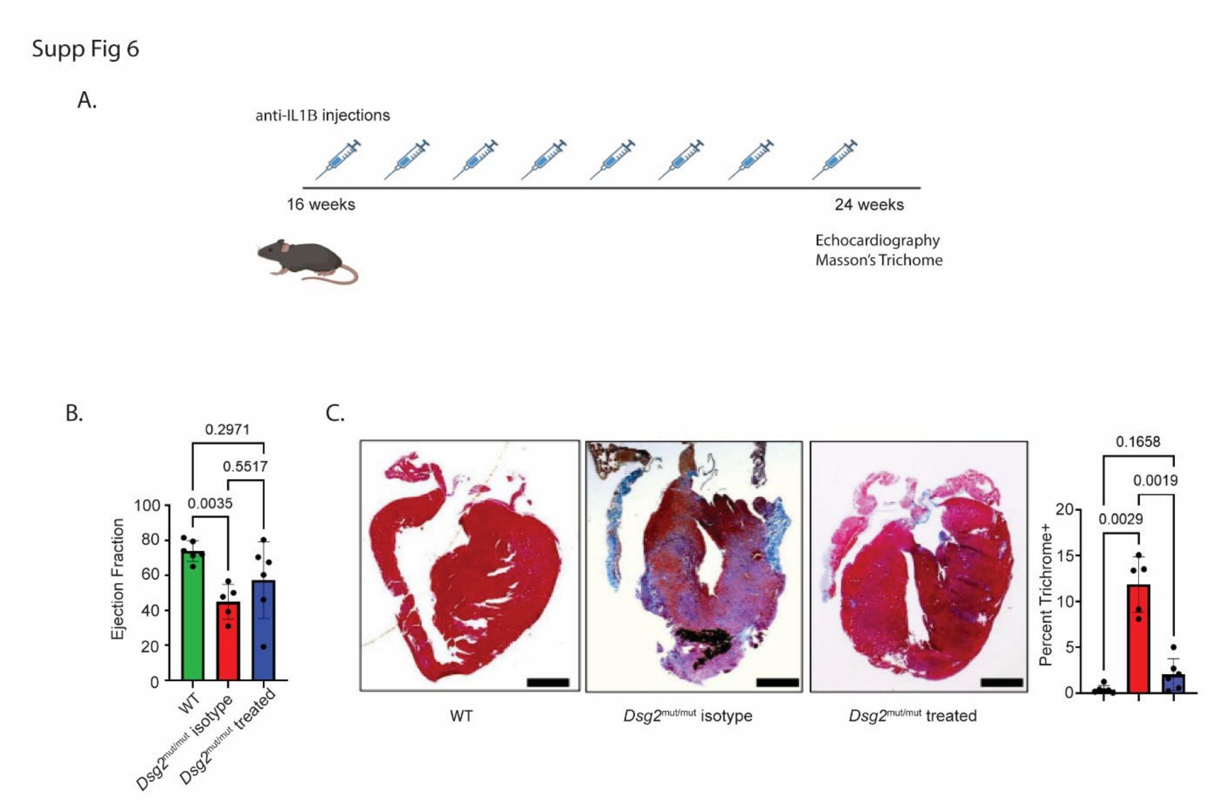


**Fig S6 Late IL1β blockade in *Dsg2*$^{mut/mut}$ mice**

(**A**) Study design outlining treatment schedule of neutralizing IL1β antibody. (**B**) Measurement of ejection fraction (n=6, 5, and 6 for each group respectively). (**C**) Representative Masson's trichrome images from each treatment group and quantification of fibrosis percentage (n=6, 5, and 6 for each group respectively). Brown-Forsythe and Welch ANOVA testing was used for graphs from (**B**) and (**C**).

Supp Fig 7

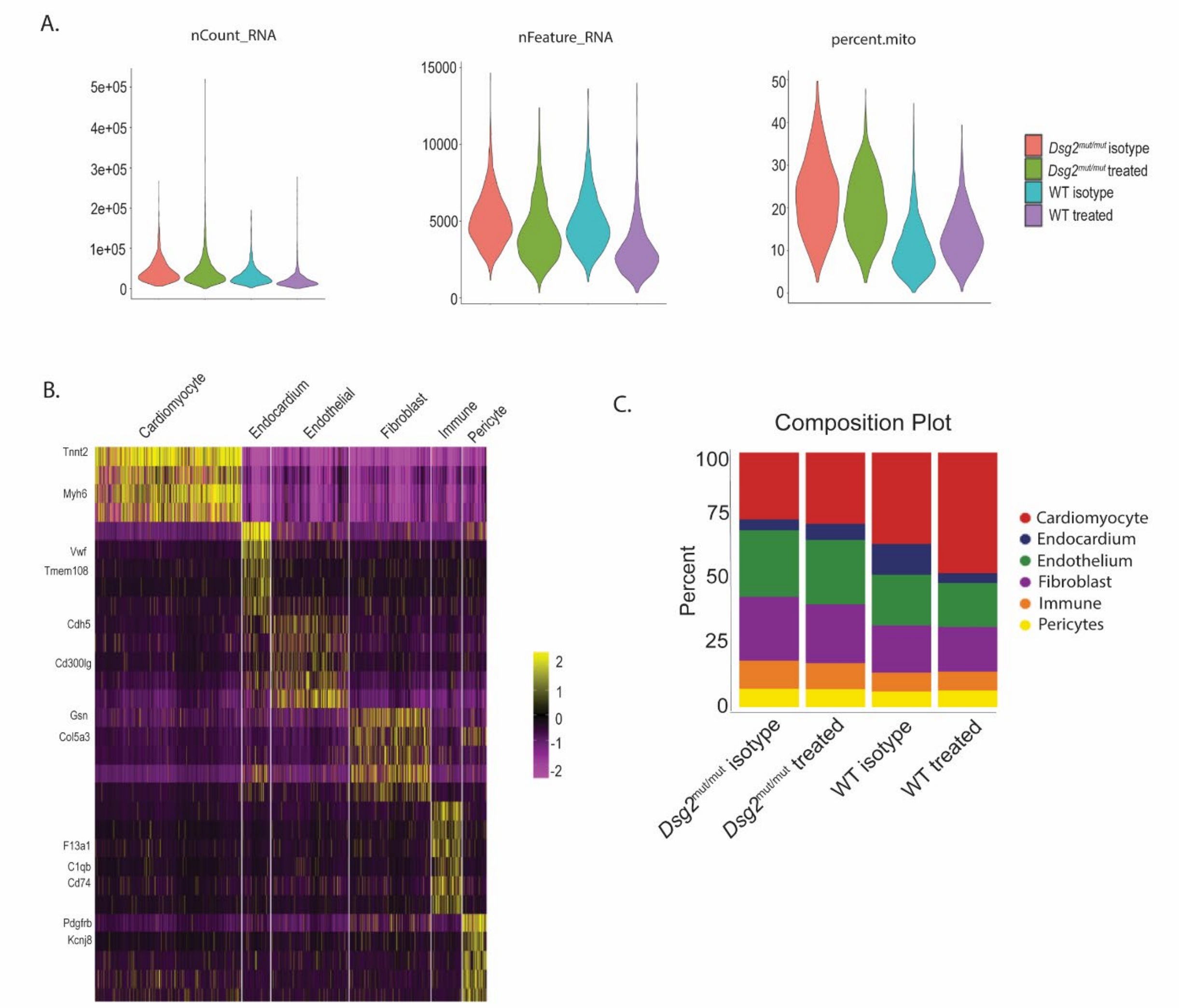


**Fig S7 Mouse single nuclei RNA sequencing with iCell8cx system**

(**A**) Major QC cutoffs for single cell RNA sequencing data. RNA count: $1000<n<300000$, Feature count: $500<n<20000$, Mitochondrial percentage: $n<30$. (**B**) Heatmap displaying major canonical gene expression across different cell types. (**C**) Composition plots comparing major cell populations across treatment groups.

| **Sample** | **Condition/Mutation** | **Gender** | **Age (years)** | **DV200** | **Used for spatial** |
|---|---|---|---|---|---|
| 11-42 | Donor control | F | 48 | 39 | |
| 11-261 | Donor control | F | 37 | 43 | X |
| 13-20 | Donor control | M | 45 | 37 | |
| 13-192 | Donor control | F | 21 | 35 | |
| 13-198 | Donor control | M | 11 | 38 | |
| 13-79 | Donor control | F | 19 | 42 | X |
| 13-96 | Donor control | M | 28 | 39 | |
| 13-152 | Donor control | M | 35 | 30 | |
| P0202 | Donor control | F | 8 | 32 | |
| P0172 | Donor control | M | 7 | 34 | |
| P0156 | Donor control | F | .75 | 36 | |
| P0159 | Donor control | M | 1 | 33 | |
| 19-14 | *PKP2* | M | 46 | 46 | X |
| 20-55 | *DSP* | F | 13 | 38 | |
| 20-283 | *DSP* | M | 14 | 44 | X |
| P0166 | *PKP2* | NA | NA | 35 | |
| 398-100 | *PKP2* | F | NA | 54 | X |
| T1077 | *DSP* | M | 61 | 32 | |

**Table S1 List of human samples used for single nuclei sequencing and spatial transcriptomics**

| **Human Gene Symbol** | **Mouse ortholog** |
|---|---|
| *AFF3* | *Aff3* |
| *SMOC2* | *Smoc2* |
| *TENM4* | *Tenm4* |
| *PDE5A* | *Pde5a* |
| *PLCB4* | *Plcb4* |
| *IL1RAPL1* | *Il1rapl1* |
| *SGIP1* | *Sgip1* |
| *PALLD* | *Palld* |
| *IGF1* | *Igf1* |
| *FAP* | *Fap* |
| *DDAH1* | *Ddah1* |
| *ASPN* | *Aspn* |
| *RUNX2* | *Runx2* |
| *MXRA5* | *Mxra5* |
| *LTBP2* | *Ltbp2* |
| *THBS4* | *Thbs4* |
| *MEOX1* | *Meox1* |
| *FN1* | *Fn1* |
| *POSTN* | *Postn* |
| *RUNX1* | *Runx1* |
| *CILP* | *Cilp* |
| *MEOX2* | *Meox2* |
| *NREP* | *Nrep* |
| *CCN2* | *Ccn2* |

**Table S2 List of human fibroblast genes (and mouse orthologs) upregulated in ACM used to build gene signature score.**

| **Human Gene Symbol** | **Mouse ortholog** |
|---|---|
| *WASHC2A* | *Washc2* |
| *NLRP3* | *Nlrp3* |
| *ZBTB16* | *Zbtb16* |
| *FOS* | *Fos* |
| *CSGALNACT1* | *Csgalnact1* |
| *SH3BP5* | *Sh3bp5* |
| *LMNA* | *Lmna* |
| *EMP1* | *Emp1* |
| *GLIS3* | *Glis3* |
| *ANXA1* | *Anxa1* |
| *ELL2* | *Ell2* |

| | |
|---|---|
| *FOSB* | *Fosb* |
| *ACSL1* | *Ascl1* |
| *SRGAP1* | *Srgap1* |
| *ARHGEF10L* | *Arhgef10l* |
| *NAMPT* | *Nampt* |
| *PAPSS2* | *Papss2* |
| *TFRC* | *Tfrc* |
| *SLC1A3* | *Slc1a3* |
| *MAN1A1* | *Man1a* |
| *CPM* | *Cpm* |
| *SRGN* | *Srgn* |
| *IGSF21* | *Igsf21* |

**Table S3 List of human myeloid genes (and mouse orthologs) upregulated in ACM used to build gene signature score.**

| Fold Change |
|---|
| ≤1 |
| 1-2 |
| 2-4 |
| 4-8 |
| 8-12 |
| **Value** |

| Cytokine | Dsg2$^{mut/mut}$ (Isotpye) vs WT (Isotype) | | Dsg2$^{mut/mut}$ (1mg/kg IL-1β) vs WT (Isotype) | |
|---|---|---|---|---|
| | Fold Change | P-Values | Fold Change | P-Values |
| CCL6 | 4-8 | P<0.001 | 1-2 | 0.874 |
| CCL22 | 2-4 | P<0.05 | ≤1 | 0.997 |
| CD14 | 4-8 | P<0.05 | 1-2 | 0.630 |
| CD40 | 8-12 | 0.108 | 2-4 | P<0.05 |
| CXCL2 | 8-12 | P<0.05 | 4-8 | 0.098 |
| CXCL9 | 2-4 | P<0.05 | 1-2 | 0.565 |
| IFN-y | 8-12 | P<0.01 | 2-4 | P<0.05 |
| IGFBP-2 | 4-8 | P<0.05 | 1-2 | 0.140 |
| IGFBP-3 | 2-4 | P<0.05 | ≤1 | 0.950 |
| IL-1β | 8-12 | P<0.01 | 2-4 | 0.354 |
| IL-3 | ≤1 | 0.938 | 2-4 | 0.246 |
| MPO | 4-8 | P<0.05 | ≤1 | 0.296 |
| Osteopontin | **19** | P<0.01 | 2-4 | 0.447 |
| Postn | 4-8 | P<0.05 | 2-4 | 0.430 |

**Table S4 Selected list of pro-inflammatory and pro-fibrotic cytokines changed following early IL1β blockade in *Dsg2$^{mut/mut}$* mice.**

Full list of genes is available in data supplement.
